# Alternative Cdc20 translational isoforms bypass the spindle assembly checkpoint to control mitotic arrest duration

**DOI:** 10.1101/2021.10.24.465552

**Authors:** Mary Jane Tsang, Iain M. Cheeseman

## Abstract

Mitotic chromosome segregation defects activate the Spindle Assembly Checkpoint (SAC), which inhibits the APC/C co-activator Cdc20 to induce a prolonged cell cycle arrest. Once errors are corrected, the SAC is silenced thereby allowing anaphase onset and mitotic exit to proceed. However, in the presence of persistent, unresolvable errors, cells can undergo “mitotic slippage”, exiting mitosis into a tetraploid G1 state and escaping the cell death that results from a prolonged arrest. The molecular logic that allows cells to balance these dueling mitotic arrest and slippage behaviors remains unclear. Here we demonstrate that human cells modulate their mitotic arrest duration through the presence of conserved, alternative Cdc20 translational isoforms. Translation initiation at downstream start sites results in truncated Cdc20 isoforms that are resistant to SAC-mediated inhibition and promote mitotic exit even in the presence of mitotic perturbations. Targeted molecular changes or naturally-occurring mutations in cancer cells that alter the relative Cdc20 isoform levels or its translational regulatory control modulate both mitotic arrest duration and anti-mitotic drug sensitivity. Our work reveals a critical role for the differential translational regulation of Cdc20 in mitotic arrest timing, with important implications for the diagnosis and treatment of human cancers.

## Full-length Cdc20 protein is not essential in human cells due to the presence of alternative Cdc20 isoforms

Cdc20 is a critical co-activator of the anaphase-promoting complex, also known as the cyclosome (APC/C), an E3 ubiquitin ligase that directs the ubiquitination and degradation of mitotic substrates to promote chromosome segregation and mitotic exit. To define the molecular mechanisms that balance the opposing roles of Cdc20 in SAC signaling and mitotic exit (Fig. 1A), we performed functional studies in human HeLa cells. Cdc20 depletion results in a well-documented mitotic arrest due to failure to activate the APC/C 2-4. Consistent with prior reports, we observed a clear mitotic arrest in HeLa cells following treatment with Cdc20 siRNAs or using an inducible CRISPR/Cas9 gene-targeting strategy 5 with an sgRNA recognizing a region within exon 3 (sgExon3) (Fig. 1B). In contrast, Cas9-mediated DNA cleavage with guides targeting a region near the start codon (sgM1) or within exon 1 (sgExon1) of the CDC20 gene did not cause a potent mitotic arrest. To define the basis for the differential effects of the Cdc20 gene disruptions, we analyzed Cdc20 protein levels by Western blotting using antibodies recognizing the C-terminus of human Cdc20 (aa 450-499). Cdc20 does not undergo alternative mRNA splicing within its coding sequence, and thus prior work has assumed that Cdc20 exists as a single protein isoform of ∼55 kDa. However, in addition to the presence of a protein matching the predicted molecular weight of full-length Cdc20 (55 kDa), this antibody detected two lower molecular-weight species. These protein bands were eliminated by Cdc20 siRNA treatment (Fig. 1C), indicating that they originate from the Cdc20 mRNA. Similar lower molecular-weight Cdc20 species were also detected in multiple human cancer cell lines (A549, DLD-1, and U2OS) and the non-transformed hTERT RPE-1 cell line (Extended Fig. 1A). These additional Cdc20 protein bands were not due to phosphorylation (Extended Fig. 1B) and were present throughout the cell cycle, starting in S phase and persisting in cells undergoing a prolonged mitotic arrest (Extended Fig. 1C).

**Figure 1.**
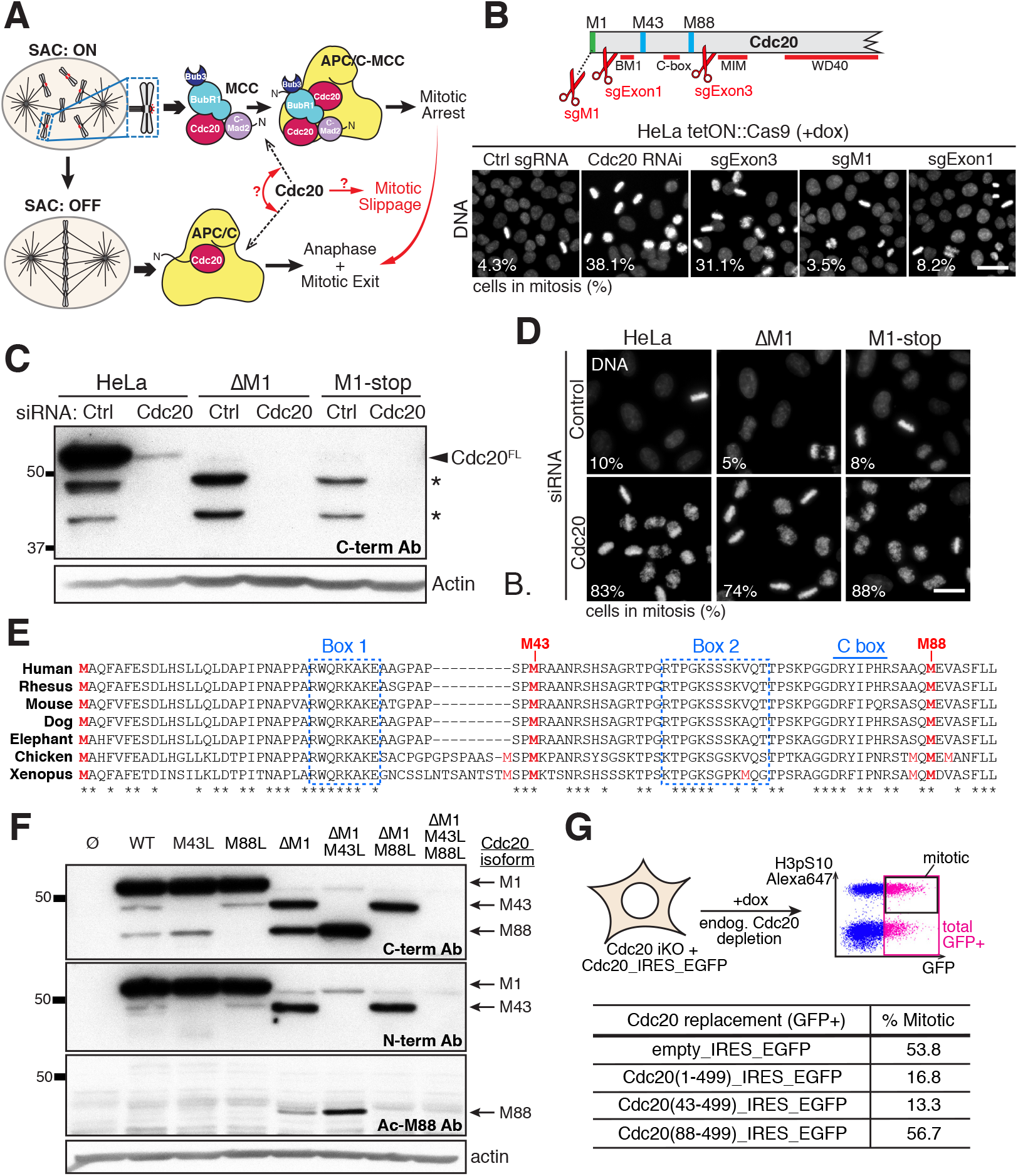
Full-length Cdc20 protein is not essential in human cells due to the presence of alternative Cdc20 translational isoforms. (A) Illustration highlighting the opposing roles of Cdc20 as the key target of the SAC and as an essential APC/C cofactor required for mitotic progression. Regulating the roles of Cdc20 in SAC signaling and mitotic exit may alter SAC efficacy and the extent of mitotic slippage (indicated in red). (B) Top, diagram of the Cdc20 open reading frame indicating the targeting of sgR-NAs and the presence of critical motifs. Bottom, representative images of DNA staining (Hoechst) to show the mitotic arrest behavior of HeLa cells following treatment with Cdc20 siRNAs or using an inducible CRISPR/Cas9 gene targeting strategy with sgRNAs recognizing different regions within CDC20 gene. Numbers indicate the average percent mitotic cells for two experimental replicates. Scale bar, 40 μm. (C) Western blot showing HeLa, ΔM1, or M1-stop cells treated with control or Cdc20 siRNAs. Endogenous Cdc20 protein was detected using antibodies recognizing the C-terminus of human Cdc20 (aa 450-499). β-actin was used as loading control. (D) Representative images of DNA staining (Hoechst) to indicate the mitotic arrest behavior of HeLa, ΔM1, and M1-stop cells treated with control or Cdc20 siRNAs. Numbers indicate the percent mitotic cells. Scale bar, 20 μm. (E) Protein alignment of the N-terminal region of human Cdc20 compared to other mammals and tetrapod species with conserved motifs indicated. Conserved amino acids are indicated with an asterisk. Methionine residues are high-lighted in red. Conserved Met1, Met43, and Met88 are bolded. (F) Western blot showing the presence of translation initiation at the Met1, Met43, and/or Met88 start codons. Wild-type and the indicated Cdc20 mutants were expressed ectopically in mitotically-enriched M1-stop cells depleted of endogenous Cdc20 protein using RNAi. β-actin was used as loading control. (G) Top, Schematic of FACS analysis of cells constitutively expressing the indicated constructs and the sgEx-on3 guide RNA to determine the fraction of mitotic cells. Endogenous Cdc20 protein was depleted with Cas9 induction and the percent of GFP-positive cells in mitosis was quantified. High levels of histone H3 phosphorylated at serine residue 10 (pS10) were used as a marker of mitosis. Bottom, numbers indicate the average percent of GFP-positive cells in mitosis for the indicated construct from two experimental replicates.

Although Cdc20 is essential for viability (6,7; Cancer Dependency Map (depmap.org)), we were able to isolate stable clonal cell lines lacking the canonical full-length Cdc20 protein (Fig. 1C). First, we isolated a homozygous mutant cell line lacking the M1 ATG start codon (ΔM1; see Extended Fig. 1D for sequence information). Second, we isolated a mutant containing insertions after the L14 residue that result in premature stop codons for both CDC20 alleles (M1-stop; see Extended Fig. 1E for sequence information). For both the ΔM1 and M1-stop mutants, the lower molecular-weight Cdc20 protein bands detected by Western blotting were now the major Cdc20 species present and these were eliminated by Cdc20 siRNA treatment (Fig. 1C). This suggests that the lower molecular-weight forms are not a result of degradation or proteolytic cleavage of the full-length Cdc20 protein, but instead reflect N-terminally truncated alternative protein isoforms. Despite the absence of full-length Cdc20, both the ΔM1 and M1-stop mutant cell lines were viable. The M1-stop mutant displayed a similar growth behavior (Extended Fig. 1F) and mitotic duration to control cells (Extended Fig. 1G; 49 min compared to 53 min for control HeLa cells). In contrast, the ΔM1 mutant displayed a modest growth defect (Extended Fig. 1F) and progressed through mitosis significantly faster than control cells (33 min) (Extended Fig. 1G), which may account for this growth defect. Importantly, both mutants are still dependent on Cdc20 for mitotic progression, as treatment with Cdc20 siRNAs resulted in a potent metaphase arrest similar to that observed in control HeLa cells (Fig. 1D). Together, these results demonstrate that human cells express multiple Cdc20 isoforms such that the canonical full-length Cdc20 protein is not strictly essential for viability or mitotic progression.

## Cdc20 isoforms are produced by alternative translation initiation at downstream in-frame start codons

We next sought to determine the nature of the alternative Cdc20 isoforms. As these isoforms share a common C-terminus based on Western blotting, we tagged the endogenous CDC20 gene locus with a C-terminal mEGFP-tag. Cdc20 contains potential downstream translation start sites at amino acid positions 43 and 88 that are conserved across mammals and diverse tetrapod species (Fig. 1E). Based on mass spectrometry analysis of Cdc20-mEGFP immunoprecipitates using either standard tryptic digests or the protease Lys-C, we identified peptides with N-terminal acetylation corresponding to translation initiation at the M1, M43, and M88 start sites (Extended Fig. 2A-C). To evaluate alternative translation initiation at these downstream start sites, we tested Cdc20 mutants using a replacement strategy combining untagged siRNA-resistant CDC20 cDNA constructs with RNAi-mediated depletion of endogenous Cdc20 (Fig. 1F). Using this gene-replacement strategy, the wild-type CDC20 cDNA construct recapitulated the Cdc20 isoform pattern observed in control HeLa cells with the presence of 3 isoforms indicating that the multiple Cdc20 isoforms can be generated from a single mRNA transcript. Importantly, mutating M43 or M88 to leucine selectively eliminated the corresponding protein products demonstrating that these start codons are responsible for the truncated Cdc20 isoforms. Deletion of the M1 start codon abrogated expression of full-length Cdc20 and also resulted in increased levels of both truncated translational isoforms. Mutating M43 or M88 to leucine in this ΔM1 construct again eliminated the expression of the M43 or M88 isoform, respectively.

**Figure 2.**
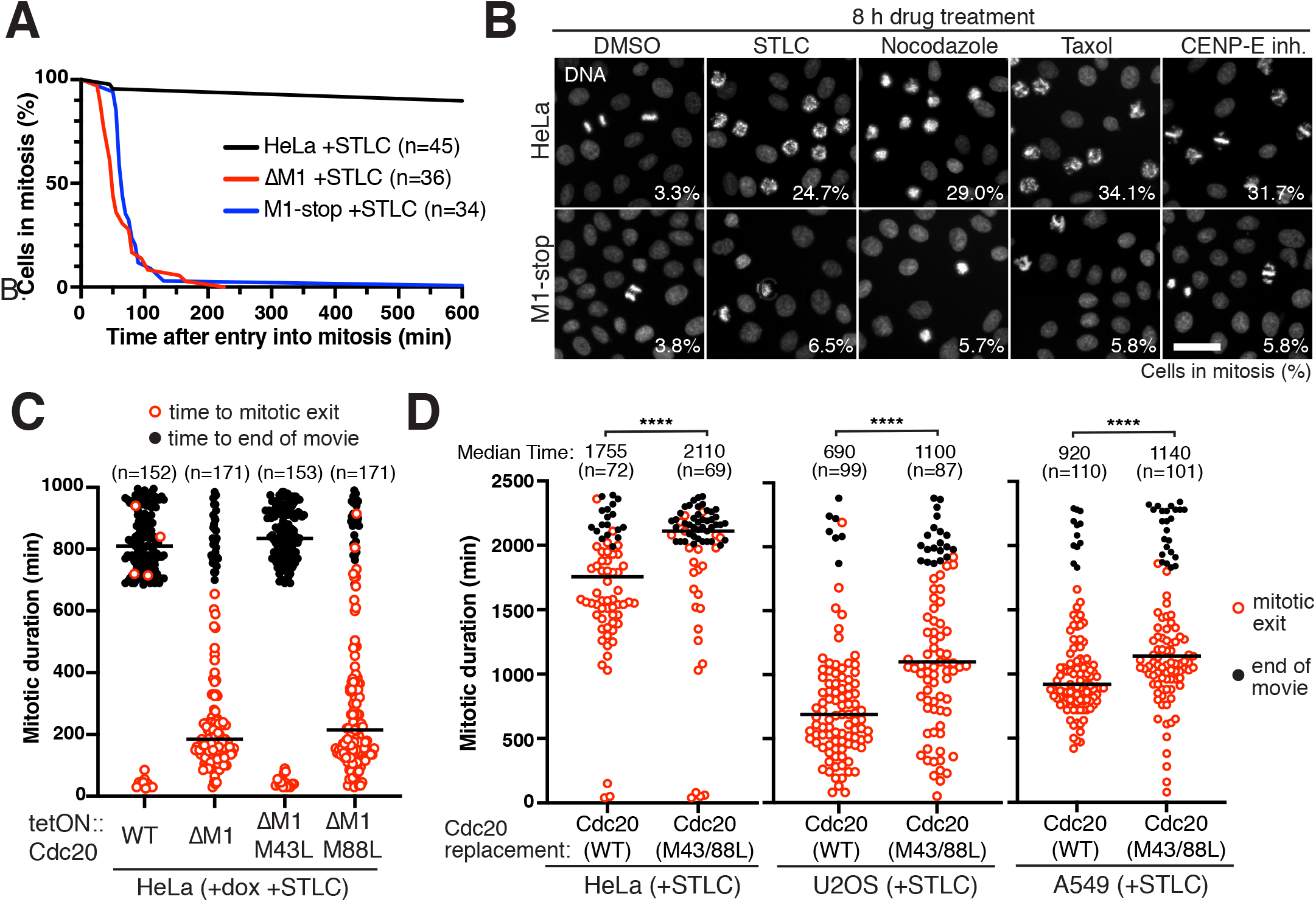
Truncated Cdc20 isoforms are inefficient targets of the SAC and promote mitotic slippage in cancer cell lines. (A) Cumulative frequency distribution for the fraction of cells in mitosis at the indicated time after entry into mitosis (mitotic arrest duration) for HeLa, ΔM1, and M1-stop cells treated with 10 μM STLC. (B) Representative images of DNA staining (Hoechst) to show the mitotic arrest behavior of HeLa or M1-stop cells treated for 8 hr with a range of anti-mitotic drugs that activate the SAC. Indicated is the average percent mitotic cells of two experimental replicates. Scale bar, 40 μm. (C) Mitotic arrest duration of individual HeLa cells treated with 10 μM STLC and 50 ng/μl doxycycline to induce expression of the indicated doxycycline-inducible CDC20 constructs. Cells entering mitosis in the first 325 min of time lapse experiments were included in analyses. Open red circles indicate cells that exit mitosis. Closed black circles indicate cells that remained arrested in mitosis till the end of the time lapse. Bars correspond to median. (D) Mitotic arrest duration in the presence of 10 μM STLC for individual HeLa, U2OS, or A549 cells with Cdc20 replacement with either wild-type CDC20 cDNA or a Cdc20 M43L M88L mutant construct. Cells entering mitosis in the first 450 min (HeLa) or 600 min (U2OS/A549) of time lapse experiments were included in analyses. Symbols are as described in (C). Indicated are the median mitotic duration times across two experimental replicates and statistics from Mann-Whitney Test (**** = p < 0.0001).

To test the functional properties of the alternative Cdc20 isoforms, we analyzed their cellular localization and ability to promote mitotic progression. Similar to full-length Cdc20, N-terminal mEGFP-Cdc20 fusions of the M43 and M88 isoforms localized to kinetochores in both untreated and nocodazole-treated cells (Extended Fig. 2D), consistent with Cdc20 kinetochore recruitment occurring through motifs in its downstream regions 8,9. Cdc20 depletion using CRISPR/ Cas9-mediated gene targeting with an sgExon3 guide RNA results in a mitotic arrest (Fig. 1B, G). This mitotic arrest phenotype was rescued by expression of guide RNA-resistant versions of full-length Cdc20 (1-499) or Cdc20 (43-499), but not Cdc20 (88-499) (Fig. 1G). Thus, both the M43 and M88 isoforms localize to kinetochores, but only the M43 isoform is able to fully complement the loss of endogenous Cdc20 protein in promoting mitotic progression. The inability of the M88 isoform to effectively promote mitotic progression is likely due to the absence of the critical “C-box” motif at residues 77-83, which is required for efficient binding to the APC/C (Fig. 1E; 10-12). Together, our results demonstrate that human cells express alternative Cdc20 translational isoforms that confer mitotic progression and viability in cells lacking full-length Cdc20.

## Truncated Cdc20 isoforms are inefficient targets of the SAC and promote mitotic slippage

Although cells lacking full-length Cdc20 are viable, we next considered whether these mutants have altered ability to arrest in mitosis in response to the Spindle Assembly Checkpoint. Both the M43 and M88 isoforms lack a conserved motif (Box1 or BM1; aa 27-34) that is required for robust Cdc20-Mad2 interactions and SAC signaling (Fig. 1E) 13-15. Treatment with the Eg5/Kif11-inhibitor STLC prevents bipolar spindle formation, resulting in potent SAC activation and an extended mitotic arrest. In time-lapse experiments, after entering mitosis, STLC-treated control HeLa cells remained arrested in mitosis for the duration of our analysis (>10 h) (Fig. 2A). In contrast, both the ΔM1 and M1-stop mutant cell lines displayed potent SAC defects as they were able to exit mitosis within a few hours, despite the presence of STLC (Fig. 2A). Similar SAC defects were also observed when M1-stop cells were treated with diverse anti-mitotic drugs (Fig. 2B).

To test whether the mitotic slippage behavior of the mutant cell lines is due to premature APC/C activation, we treated cells with both STLC and the APC/C-inhibitor proTAME 16. APC/C inhibition suppressed the premature mitotic exit observed in ΔM1 and M1-stop cells, resulting in a prolonged mitotic arrest (Extended Fig. 3A). In addition to Cdc20, Cdh1 acts as a co-activator of the APC/C in late mitosis 17, suggesting that Cdh1 could substitute for Cdc20 in the mutant cell lines to promote premature APC/C activation. However, Cdh1 depletion by RNAi did not alter the premature mitotic exit of the ΔM1 and M1-stop mutant cell lines (Extended Fig. 3B). Finally, Cdc20 replacement using the wild-type CDC20 cDNA restored prolonged mitotic arrest behavior to both the

**Figure 3.**
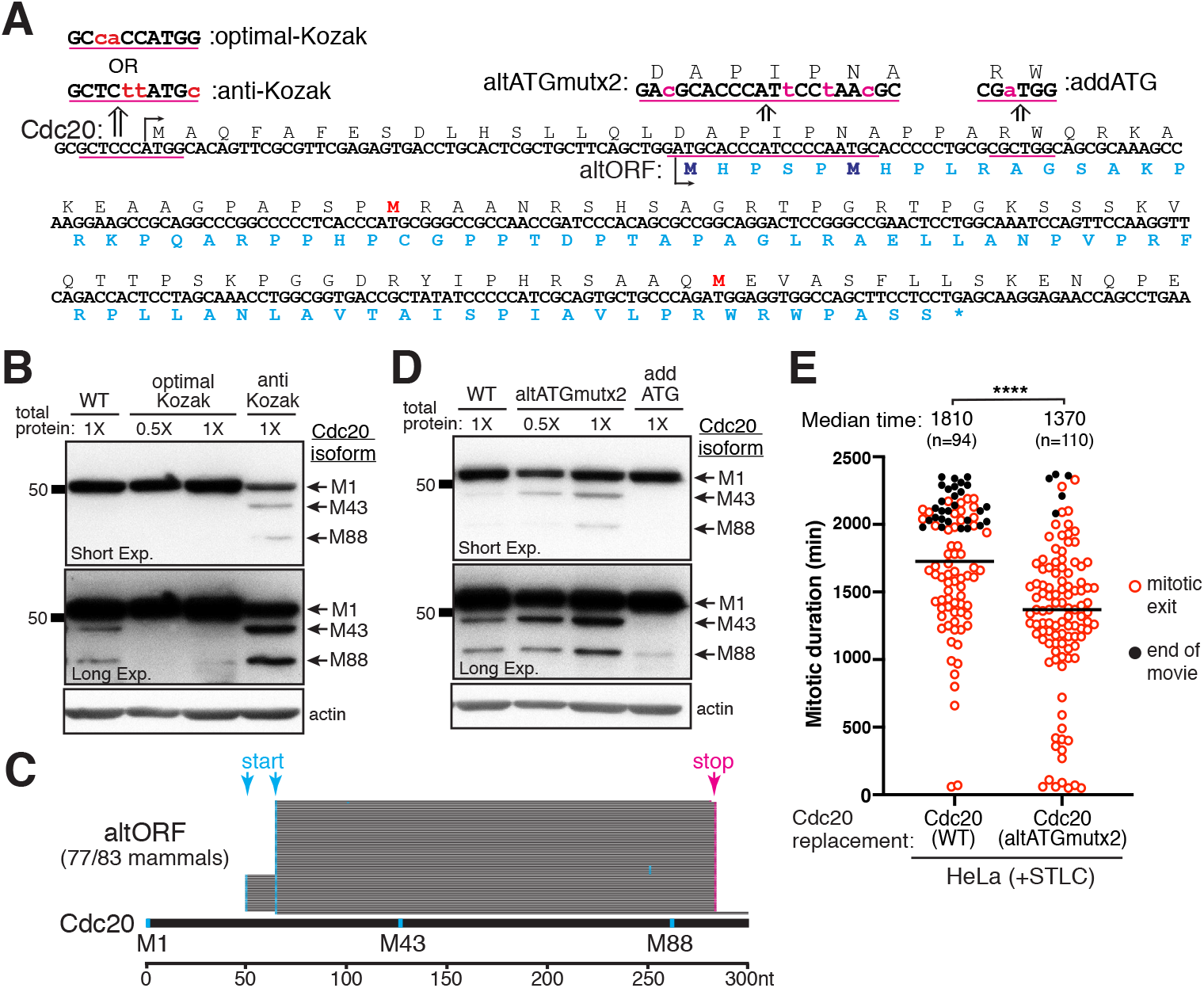
Leaky scanning and translation initiation at alternative out-of-frame start codons modulate Cdc20 isoform expression levels. (A) Analysis of human CDC20 nucleic acid sequence reveals two alternative out-of-frame start codons between Met1 and Met43. The amino acid sequence of the predicted alternative open reading frame (altORF) is indicated in cyan, with the methionines bolded. Mutations to alter the translation-initiation context of Met1 to either a consensus Kozak sequence (optimal-Kozak) or an antiKozak sequence (anti-Kozak) are underlined and highlighted in magenta. Targeted silent mutations to disrupt the out-of-frame start codons (altATGmutx2) or introduce an additional out-of-frame start codon before the Met43 start site (addATG) are similarly indicated. (B) Western blot showing CDC20 constructs with mutations to alter the translation-initiation context of Met1 to either a consensus Kozak sequence (optimal-Kozak) or an antiKozak sequence (anti-Kozak). Constructs were expressed in mitotically-enriched M1-stop cells depleted of endogenous Cdc20 protein. The translation products were detected using antibodies recognizing the human Cdc20 C-terminus (aa 450-499). β-actin was used as loading control. (C) Conservation analysis for the presence of Cdc20 altORF across various mammalian species. The altORF was detected in 77 out of 83 species analyzed. For each species, start and stop codons are mapped relative to human Cdc20 protein sequence and indicated in cyan and magenta respectively. (D) Similar Western blot as in (B) except showing CDC20 constructs with targeted silent mutations to disrupt the out-of-frame start codons (altATGmutx2) or introduce an additional out-of-frame start codon before the Met43 start site (addATG). (E) Mitotic arrest duration in the presence of 10 μM STLC for individual HeLa cells in which the endogenous Cdc20 protein is replaced with either wild-type siRNA-resistant CDC20 cDNA or the altAT-Gmutx2 mutant construct. Cells entering mitosis in the first 450 min of time lapse experiments were included in analyses. Open red circles indicate cells that exit mitosis. Closed black circles indicate cells that remained arrested in mitosis till the end of the time lapse. Bars correspond to median. Indicated are the median mitotic duration times across two experimental replicates and statistics from Mann-Whitney Test (**** = p < 0.0001).

ΔM1 and M1-stop cell lines (Extended Fig. 3C). In addition, similar SAC defects were observed following the acute depletion of full-length Cdc20 using Cas9 induction in cell lines expressing either the sgM1 or sgExon1 guide RNAs (Extended Fig. 3D). Therefore, the SAC defects in these mutant cell lines are due to the absence of full-length Cdc20 rather than other potential second-site mutations or long-term adaptation in the stable mutant cell lines. Together, these results demonstrate that the loss of full-length Cdc20 impairs SAC function and results in an APC/C- and Cdc20-dependent mitotic exit in the presence of anti-mitotic drugs.

The premature mitotic exit observed in the ΔM1 and M1-stop mutant cell lines could reflect a defect in the upstream SAC signaling pathway. To assess whether SAC activation occurs in these mutants, we tested the localization of the SAC proteins Bub1 and Mad2, which are recruited to unattached kinetochores to trigger checkpoint signaling 18. Bub1 and Mad2 localized to kinetochores in ΔM1 and M1-stop mutant cell lines treated with the microtubule-depolymerizing drug nocodazole similar to control HeLa cells (Extended Fig. 3E). However, despite evidence for upstream SAC activation, both the ΔM1 and M1-stop mutant cell lines behaved functionally as if the SAC was absent (Fig. 2A). Indeed, the premature mitotic exit observed for the ΔM1 mutant in the presence of STLC was not exacerbated further by weakening the SAC by treatment with either the Mps1 inhibitor, AZ3146, which targets the most upstream component of the SAC signaling cascade (Extended Fig. 3F, Extended Table 1), or using siRNAs against Mad2, a key component of the SAC effector complex (Extended Fig. 3G, Extended Table 1). However, targeted SAC inhibition using AZ3146 or Mad2 siRNAs was able to further reduce the arrest of the M1-stop mutant, resulting in a mitotic duration similar to that of the ΔM1 mutant (Extended Table 1). Together, our results suggest that the truncated Cdc20 isoforms in the ΔM1 and M1-stop cell lines are not effectively targeted and inhibited by the SAC, resulting in premature APC/C activation and mitotic slippage in the presence of anti-mitotic drugs.

## Cdc20 translational isoforms modulate mitotic arrest duration

In the absence of full-length Cdc20, the truncated Cdc20 isoforms are not effectively inhibited by the SAC, resulting in premature APC/C activation and mitotic exit. However, control cells co-express multiple Cdc20 translational isoforms, including the full-length protein (Fig. 1C; Extended Fig. 1A), such that these isoforms must compete for APC/C binding. To test whether the truncated Cdc20 isoforms can modulate mitotic arrest timing in the presence of full-length Cdc20, we altered the relative Cdc20 isoform levels using ectopic expression of CDC20 cDNA constructs. Overexpression of wild-type CDC20 encoding all three isoforms did not substantially alter the mitotic arrest duration in the presence of STLC (Fig. 2C). In contrast, expression of a CDC20-ΔM1 construct to increase the levels of the shorter Cdc20 isoforms relative to full-length Cdc20 (Extended Fig. 4A) caused a dramatic increase in mitotic slippage with cells exiting mitotic arrest within a few hours (Fig. 2C). Preventing the increased expression of the M43 isoform in the CDC20-ΔM1 construct using an M43L mutation fully suppressed the induced mitotic slippage phenotype, whereas an M88L mutation had little effect (Fig. 2C). Thus, increased levels of the Cdc20 (43-499) isoform are sufficient to promote mitotic slippage even in the presence of full-length Cdc20 protein.

**Figure 4.**
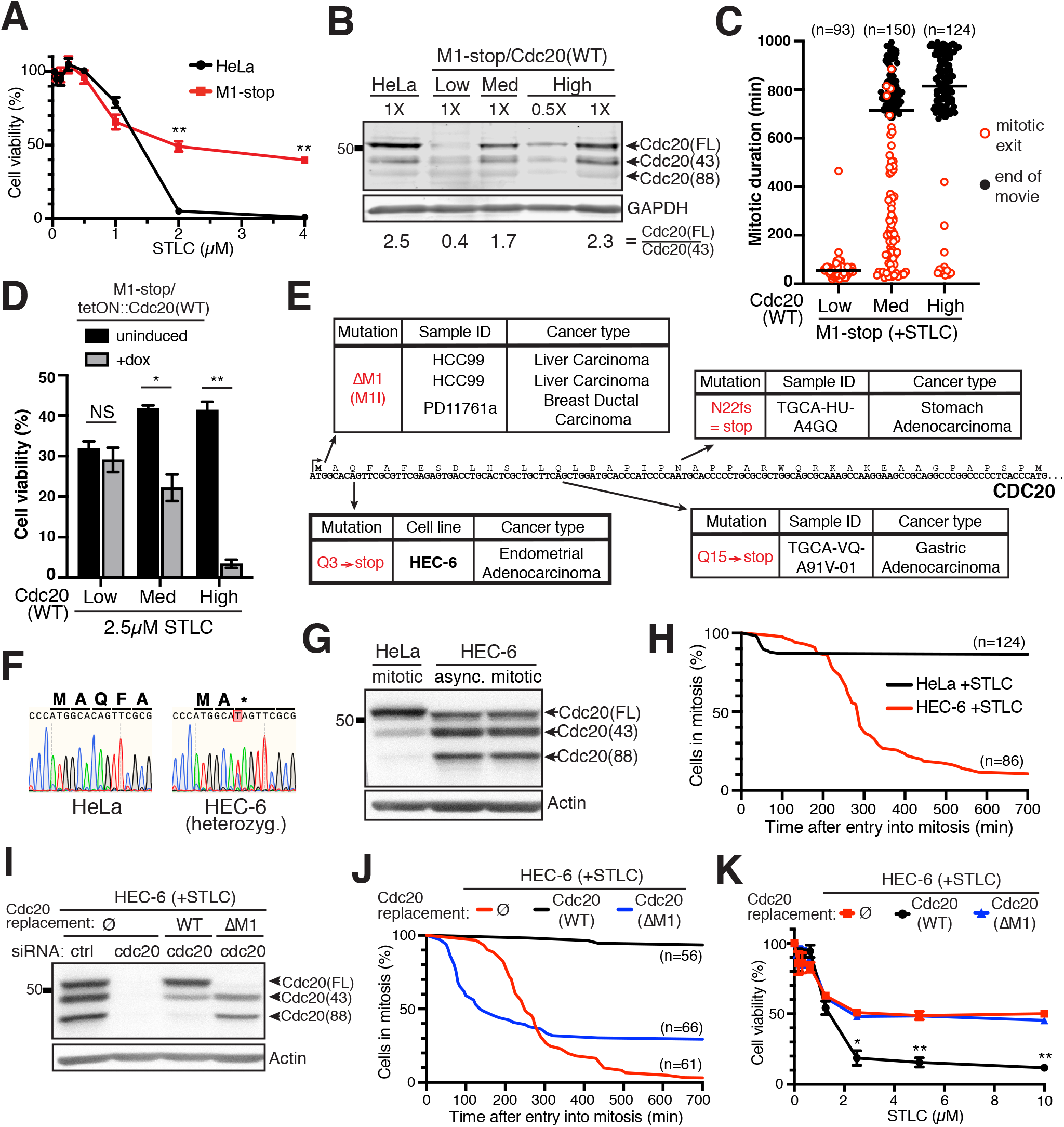
Cdc20 translational isoform levels alter cancer cell anti-mitotic drug sensitivity. (A) Sensitivity of HeLa or M1-stop cells to increasing concentrations of the anti-mitotic drug STLC. Cell viability was determined by MTT assay in triplicate following 72 h drug treatment. Error bars indicate SEM of three experimental replicates. Statistics from Student’s two-sample t-Test with two-tailed distribution comparing HeLa and M1-stop cell viabilities per drug concentration (** = p < 0.01). (B) Western blot showing representative mitotically-enriched control HeLa or representative clones of M1-stop mutant with low, medium, or high expression of the doxycycline-inducible wild-type CDC20 construct induced with 20 ng/μl doxycycline. Cdc20 was detected using antibodies recognizing the N-terminus of human Cdc20 (aa 1-175) and the ratio of full-length protein to Cdc20(43-499) was quantified. Indicated is the average of three experimental replicates GAPDH was used as loading control. (C) Mitotic arrest duration in the presence of 10 μM STLC for the representative clones in (B) treated with 20 ng/ul doxycycline. Open red circles indicate cells that exit mitosis. Closed black circles indicate cells that remained arrested in mitosis till the end of the time lapse. Bars correspond to median. (D) Cell viability at the indicated STLC concentration for the representative clones in (B-C) without induction or induced with 20 ng/μl doxycycline. Cell viability was determined by MTT assay in triplicate following 72 hr drug treatment. Error bars indicate SEM of three experimental replicates. Statistics from Student’s two-sample t-Test with two-tailed distribution (* = p < 0.05, ** = p < 0.01). (E) Survey of tumors and cancer cell lines using public databases reveals multiple distinct genetic mutations within CDC20 that are predicted to selectively deplete the full-length Cdc20 protein. Indicated are the genetic change, the sample ID or cell line containing the mutation, and the cancer type. (F) Sanger sequencing of CDC20 gene in HEC-6 cell line with the M1 start codon region highlighted. (G) Western blot of control HeLa cells compared to HEC-6 cells. Cdc20 protein was detected using antibodies recognizing the human Cdc20 C-terminus (aa 450-499). β-actin was used as loading control. (H) Cumulative frequency distribution for the fraction of cells in mitosis at the indicated time after entry into mitosis (mitotic arrest duration) for HeLa and HEC-6 cells treated with 10 μM STLC. (G) Similar Western blot as in (G) except HEC-6 cells expressed mEGFP alone or the indicated CDC20 construct and were treated with either control or Cdc20 siRNAs. (J) Similar cumulative frequency distribution as in (H) except HEC-6 cells expressed mEGFP alone or had the endogenous Cdc20 protein replaced with the indicated CDC20 construct. (K) Similar STLC drug sensitivity as in (A) except for HEC-6 cells expressing mEGFP alone or with the endogenous Cdc20 protein replaced with the indicated CDC20 construct. Cell viability was determined by MTT assay following 92 hr drug treatment. Error bars indicate SEM of three experimental replicates. Statistics from Student’s two-sample t-Test with two-tailed distribution comparing the wild-type CDC20 replacement cell viabilities to either control HEC-6 or CDC20-ΔM1 replacement per drug concentration (* = p < 0.05, ** = p < 0.01).

To evaluate the consequences of eliminating the truncated Cdc20 isoforms, we next analyzed mitotic arrest timing in HeLa cells, the osteosarcoma U2OS cell line, and the lung adenocarcinoma A549 cell line. All three cell lines express the truncated Cdc20 isoforms (Fig. 1F, Extended Fig. 4B) and undergo mitotic slippage due to Cdc20-mediated APC/C activation (Extended Fig. 4C). RNAi-based replacements with a wild-type CDC20 cDNA construct recapitulated the mitotic durations observed in the corresponding control cell lines, with median arrest times of 1755 min, 690 min, and 920 min for HeLa, U2OS, and A549, respectively (Fig. 2D, Extended Fig. 4C). Strikingly, replacement of endogenous Cdc20 with a Cdc20 M43L M88L mutant to prevent expression of the truncated isoforms resulted in a significant increase in the mitotic arrest duration for each cell line, with the median arrest times increasing to 2110 min, 1100 min, and 1140 min for HeLa, U2OS, and A549, respectively (Fig. 2D). Overall, these results indicate that the truncated Cdc20 isoforms contribute to mitotic slippage in diverse cancer cell lines and our results support a model in which the relative levels of Cdc20 isoforms influence the mitotic arrest behavior of individual cells.

## Leaky scanning and translation initiation at alternative out-of-frame start codons modulate Cdc20 isoform expression levels

Given our observation that the relative levels of the different Cdc20 translational isoforms impact mitotic arrest duration, we next investigated the molecular mechanisms that control Cdc20 translation start site selection. Eukaryotic translation initiation is typically accomplished by a scanning mechanism in which the 40S ribosomal subunit is loaded at the 5’-cap and translocates along the mRNA until it initiates translation at the first AUG encountered 19. However, translation at down-stream start codons can occur by either translational re-initiation or leaky scanning in which a fraction of 40S ribosomal subunits continue scanning beyond the first AUG to initiate at a downstream AUG 20,21. Consistent with leaky ribosome scanning, deletion of the CDC20 M1 start codon in either the ΔM1 mutant cell line (Fig. 1C) or the ΔM1 ectopic construct (Fig. 1F) results in increased initiation at the downstream M43 and M88 start codons. Leaky ribosomal scanning occurs when the translational context of the first AUG is suboptimal 21. Introducing a strong consensus Kozak sequence at the M1 start site (Fig. 3A) should increase translation initiation at this start codon and reduce downstream translation initiation. Indeed, in RNAi-based replacement assays with this “optimal-Kozak” construct, we observed increased levels of the full-length Cdc20 protein compared to the wild-type construct and a concomitant decrease in translation of the M43 and M88 isoforms (Fig. 3B). Conversely, further weakening the translational context at the M1 start site with an “anti-Kozak” sequence, which still retains the AUG codon, reduced expression of the full-length protein while allowing increased translation initiation at the M43 and M88 start sites. These results are consistent with downstream translation initiation relying on leaky ribosome scanning such that the translational context surrounding the M1 start site determines the relative expression of the truncated isoforms. Importantly, this behavior would also allow the levels or activities of translation initiation factors to influence Cdc20 start site selection.

In analyzing the CDC20 mRNA sequence, we found that M43 and M88 are not the only potential start codons downstream of the annotated M1 start site. In fact, we identified two alternative out-of-frame start codons between M1 and M43 (Fig. 3A), which would be predicted to capture scanning ribosomes, preventing a subset of 40S ribosomal subunits from reaching M43 or M88. These out-of-frame start codons are conserved across mammals with a conserved stop site present downstream of the M88 codon (Fig. 3C). Translation of this alternative open reading frame (altORF) would therefore prevent initiation at the M43 and M88 start codons. Using frame shift mutant cell lines that connect the predicted altORF sequence with downstream regions of the Cdc20 protein (Extended Fig. 5A, see Extended Fig. 5B for sequence information), we were able to detect chimeric altATG-Cdc20 proteins by Western blotting (Extended Fig. 5C). This suggests that the alternative out-of-frame AUGs within CDC20 are functional for translation initiation. The presence of the alternative start sites within CDC20 is predicted to negatively impact translation initiation at the downstream M43 and M88 sites. Indeed, using our RNAi-mediated replacement strategy, we found that mutating the alternative start sites with silent mutations that do not disrupt the Cdc20 coding sequence (altATGmutx2; Fig. 3A) resulted in increased M43 and M88 isoform levels compared to wild-type controls without altering full-length protein levels (Fig. 3D). Reciprocally, introducing additional out-of-frame start codon(s) before the M43 start site (“addATG”; Fig. 3A) prevented translation of the M43 isoform, although the M88 isoform was only partially affected (Fig. 3D). Finally, we tested the contribution of these alternative out-of-frame start codons to the mitotic arrest behavior of HeLa cells. Upon STLC treatment, cells with RNAi-based replacement using the wild-type CDC20 cDNA displayed a median arrest duration of 1810 min (Fig. 3E). In contrast, replacement with the altATGmutx2 construct, which increases the relative levels of the truncated Cdc20 isoforms, showed a significant reduction in the mitotic arrest duration, with a median arrest duration of 1370 min. Thus, increasing the truncated Cdc20 isoform levels by disrupting altORF expression decreases the mitotic arrest duration (Fig. 3E) even though full-length protein levels are not substantially altered (Fig. 3D). Overall, our work suggests that expression of the M43 and M88 isoforms relies on leaky ribosome scanning downstream of the M1 start site. The presence of alternative translation sites before the M43 start codon provides a conserved mechanism to modulate translation initiation at these downstream in-frame start sites and thus affect mitotic arrest duration.

## Cdc20 translational isoform levels alter cancer cell anti-mitotic drug sensitivity

Anti-mitotic drugs, such as paclitaxel (taxol), other taxanes, and vinca alkaloids, are widely used as first-line chemotherapeutics for the treatment of breast cancer, ovarian cancer, and other cancer types 22,23. However, human cancer cells vary widely in their responses to anti-mitotic chemotherapeutics and their ability to escape from the prolonged mitotic arrest induced by treatment with these compounds 24,25. This mitotic arrest behavior has important implications for the efficacy of anti-mitotic drugs 26, with prior work suggesting that mitotic slippage limits the ability of anti-mitotic drugs to kill cancer cells 26,27. Indeed, we observed that the clonal M1-stop mutant cell line, which undergoes rapid mitotic slippage in the presence of STLC (Fig. 2A), displayed increased cell viability relative to control cells when treated with a range of anti-mitotic drugs (Fig. 4A; Extended Fig. 6A).

As the relative level of Cdc20 isoforms influences mitotic arrest duration, we sought to test the effect of these changes on anti-mitotic drug sensitivity. Control cells display a ratio of 2.5:1 for the full-length Cdc20 protein relative to the M43 isoform (Fig. 4B). To modulate the Cdc20 translational isoform ratios across a range of levels, we generated clonal cell lines expressing varying levels of the wild-type CDC20 cDNA from an inducible construct in the M1-stop mutant cell line, which lacks full-length Cdc20 (Extended Fig. 6B-C). Our analysis revealed a correlation between the relative Cdc20 isoform ratio, mitotic arrest duration, and anti-mitotic drug sensitivity (Fig 4B-D, Extended Fig 6D-E). Cell lines with low ectopic expression of full-length Cdc20 (with a ratio of 0.4:1 for full-length:M43; Fig. 4B) displayed a short mitotic arrest duration (Fig. 4C) and similar drug sensitivity to the uninduced cell line (Fig. 4D, Extended Fig. 6E) or the M1-stop mutant (Fig. 4A). In contrast, cell lines expressing a high ratio of full-length Cdc20 (2.3:1) remained arrested in mitosis for the duration of our analysis and showed significantly increased STLC sensitivity compared to uninduced controls. Interestingly, clones expressing an intermediate ratio of full-length Cdc20 protein (1.7:1) displayed heterogeneous mitotic arrest behavior (Fig. 4C). At this intermediate level of full-length Cdc20 protein, stochastic differences in Cdc20 expression in individual cells within the population may lead to variations in the Cdc20 isoform ratio that impact mitotic arrest timing. Treatment with Cdc20 siRNAs to deplete the truncated isoforms expressed from the endogenous M1-stop mutant locus suppressed this heterogeneous mitotic arrest behavior and abrogated the observed mitotic slippage (Extended Fig. 6D), but did not affect cell cycle progression in the absence of STLC (data not shown). Importantly, the heterogeneous mitotic arrest behavior observed in cells with medium Cdc20 isoform ratio correlates with an intermediate increase in anti-mitotic drug sensitivity (Fig. 4D). Together, our results indicate that, within a given cell line, the relative levels of Cdc20 isoforms modulate both mitotic arrest timing and anti-mitotic drug sensitivity.

We next considered whether genetic changes to the CDC20 coding region in cancers could impact the relative Cdc20 isoform levels. Through a survey of tumors and cancer cell lines using public databases, we identified multiple distinct genetic mutations within CDC20 that are predicted to eliminate the full-length Cdc20 protein, thus increasing the relative levels of the truncated isoforms (Fig. 4E). To evaluate the consequences of these cancer mutations, we obtained the endometrial adenocarcinoma HEC-6 cell line. Sequencing of the CDC20 locus confirmed the reported nonsense mutation (Q3-stop), but also revealed the presence of a wild-type CDC20 allele, indicating that the HEC-6 cell line is heterozygous for this CDC20 mutation (Fig. 4F). As predicted from this mutation, the HEC-6 cell line displayed a substantial reduction in the relative levels of full-length Cdc20, with a corresponding increase in the M43 and M88 isoforms (Fig. 4G). In addition, the HEC-6 cell line showed substantially reduced mitotic arrest duration in the presence of STLC relative to HeLa cells, with a median arrest duration of 285 min (Fig. 4H). To test whether this reduced mitotic arrest duration is due to the altered Cdc20 isoform levels, we performed RNAi-based replacement assays. Replacement with wild-type CDC20 cDNA resulted in increased relative levels of full-length Cdc20 (Fig. 4I) and a concomitant increase in the mitotic arrest duration, with cells remaining arrested in mitosis for >700 min in the presence of STLC (Fig. 4J). In contrast, replacement with a ΔM1 mutant abrogated expression of the full-length Cdc20 protein and resulted in a further reduction in the mitotic arrest duration, with a median arrest time of 140 min. Importantly, the relative Cdc20 isoform ratio also correlated with anti-mitotic drug sensitivity for the HEC-6 cell line. Control HEC-6 cells or cells expressing only CDC20-ΔM1 mutant cDNA were more resistant to high concentrations of STLC compared to HeLa cells (Fig. 4A, 4K). However, replacement with the wild-type CDC20 cDNA resulted in significantly increased drug sensitivity (Fig. 4K), demonstrating that drug resistance can be reversed by restoring normal Cdc20 isoform ratios.

Together, our findings reveal a mechanism to control mitotic arrest duration based on the presence of alternative Cdc20 translational isoforms that are resistant to SAC-mediated inhibition, providing a “release value” that allows cells to promote mitotic exit even in the presence of persistent errors. Genetic mutations or translational differences that influence the relative Cdc20 isoform levels would affect cancer behavior and impact anti-mitotic drug sensitivity, providing potential avenues for the diagnosis and treatment of human cancers.

## Methods

### Cell culture

HeLa, hTERT-RPE1, U2OS, and A549 cell lines were cultured in Dulbecco’s modified Eagle medium (DMEM) supplemented with 10% fetal bovine serum (FBS), 100 U/mL penicillin and streptomycin, and 2 mM L-glutamine at 37°C with 5% CO2. HEC-6 cell line was cultured in DMEM supplemented with 15% FBS and 2mM L-glutamine. Doxycycline-inducible cell lines were cultured in medium containing FBS certified tetracycline-free. spCas9 expression in inducible CRISPR/ Cas9 cell lines was induced with 1 μg/ml doxycycline hyclate (Sigma) at 24 hr intervals for 2 days. All other doxycycline-inducible constructs were induced with 10 ng/ml doxycycline hyclate, unless indicated in figure legend. Other drugs used on human cells were Nocodazole (Sigma, 330 nM), S-trityl-L-cysteine (STLC, Sigma, 10 μM), Taxol (Paclitaxel, Invitrogen, 1 μM), GSK923295 (CENP-E inhibitor, Selleck Chemicals, 100 nM), proTAME (APC/Ci, R&D Systems, 12 μM), AZ-3146 (Mps1i, Selleck Chemicals, 4 μM) unless concentration indicated otherwise in figure legend. Cells were enriched in mitosis with treatment with 330 nM nocodazole for 16-17 hrs or if indicated, 10μM STLC for 18 hrs. Cell lines were regularly tested for mycoplasma contamination.

### Cell line generation

The cell lines used in this study are described in Extended Table 3. The inducible CRISPR/Cas9 HeLa cell line was previously generated by transposition as described 5. A control sgRNA (Ctrl sgRNA, GCCGATGGTGAAGTGGTA-AG) or sgRNAs targeting different regions within the CDC20 gene (sgM1, TCGAACGCGAACTGTGCCAT; sgExon1, CCTGCACTCGCTGCTTCAGC; sgExon3, CCAGGAA-CATCAGAAAGCCT) were cloned into the sgOpti plasmid (puro-resistant, Addgene #85681) and introduced into the inducible CRISPR/Cas9 HeLa cell line by lentiviral transduction 5. Cells were selected with 0.5 μg/ml puromycin (Gibco) for 5 days.

Stable clonal cell lines lacking the canonical full-length Cdc20 protein were obtained by transfecting HeLa cells with pX330-based plasmids 28 expressing spCas9 and either the sgM1 (for ΔM1 mutant) or the sgExon1 guide RNA (for M1-stop mutant). pX330-based plasmids were transfected using X-tremeGENE-9 (Roche) together with a mCherry-expressing plasmid. Single mCherry-positive cells were fluorescence activated cell-sorted into 96-well plates. Clones were screened for successful gene editing and the nucleotide sequence of the CDC20 alleles were determined by next-generation sequencing (Genewiz Amplicon-EZ).

pBABE derivatives containing empty IRES_EGFP or different Cdc20_IRES2_EGFP constructs were transfected with Effectene (Qiagen) along with VSVG packaging plasmid into 293-GP cells for generation of retrovirus as described 29. Supernatant containing retrovirus was sterile-filtered, supplemented with 20 μg/mL polybrene (Millipore) and used to transduce inducible CRISPR/Cas9 HeLa cells expressing the sgExon3 guide RNA. At two days post-transduction, cells were selected with 375 μg/ml hygromycin (Invitrogen) for 10-14 days.

Doxycycline-inducible cell lines were generated by homology-directed insertion into the AAVS1 “safe-harbor” locus. Donor plasmid containing selection marker, the tetracycline-responsive promoter, the transgene as indicated, and reverse tetracycline-controlled transactivator flanked by AAVS1 homology arms 30 was transfected into the indicated cell line using Effectene (Qiagen) according to the manufacturer’s protocol with a pX330-based plasmid 28 expressing both sp-Cas9 and a guide RNA specific for the AAVS1 locus (GGGGCCACTAGGGACAGGAT). At two days post-transduction, cells were selected with the indicated concentration of hygromycin (375 μg/ml for HeLa and its derivatives, 100 μg/ml for U2OS, and 300 μg/ml for A549) for 10-14 days.

Lentiviral plasmids containing mEGFP only or CDC20(WT) or CDC20-ΔM1 under the control of the UbC promoter were introduced into the HEC-6 cell line by lentiviral transduction. Lentivirus was removed from cells after 4 hrs. At two days post-transduction, cells were selected with 0.25 μg/ml puromycin (Gibco) for 4 days.

### RNAi treatment and gene replacements

Custom siRNAs against Cdc20 (5’-CGGAAGACCUGC-CGUUACAUU), Cdh1 (5’-UGAGAAGUCUCCCAGUCAG), and Mad2 (5’-UACGGACUCACCUUGCUUGUU) and a non-targeting control pool (D-001206-13) were obtained from Dharmacon. siRNAs were applied at a final concentration of 50 nM, unless indicated in the figure legend. 2.5 μl Lipofectamine RNAiMAX (Invitrogen) was used per ml of final transfection medium. For gene replacements, transfection medium also contained the appropriate concentration of doxycycline hyclate to express the ectopic inducible construct. Transfection medium was changed after 6 hrs for time-lapse microscopy analyses and either 6 hrs or 20 hrs for Western blot analyses.

### Immunofluorescence microscopy

Cells for immunofluorescence were seeded on poly-L-lysine (Sigma) coated coverslips and treated with 330 nM nocodazole for 1.5 hr before pre-extraction and fixation. Cells were pre-extracted in PBS + 0.2% Triton X-100 for 1 min at 37°C before fixation with PBS + 0.2% Triton X-100 + 4% formaldehyde at room temperature for 10 min. Coverslips were washed with PBS + 0.1% Triton X-100 and blocked in Abdil (20 mM Tris-HCl pH 7.5, 150 mM NaCl, 0.1% Triton X-100, 3% BSA, 0.1% NaN3) for 30 min. Immunostaining was performed by incubating coverslips with primary antibodies diluted in Abdil for 45 min at room temperature followed by 3 consecutive washes with PBS + 0.1% Triton X-100. Cy2- and Cy5-conjugated secondary antibodies (Jackson ImmunoResearch Laboratories) were diluted 1:500 in Abdil together with 0.3 μg/ml Hoechst-33342 (Invitrogen) and incubated with coverslips for 45 min. After washing with PBS + 0.1% Triton X-100, coverslips were mounted with ProLong Gold Antifade (Invitrogen) and allowed to cure overnight before imaging.

The following primary antibodies were used: anti-centromere antibodies, ACA (1:200, Antibodies Inc, #15-234), Mad2 (1:1,000, Kops Lab 31), Bub1 (1:200, Abcam, ab54893). Images were acquired on a DeltaVision Core deconvolution microscope (Applied Precision) equipped with a CoolSnap HQ2 charge-coupled device camera and deconvolved where appropriate using the Softworx software. Z-sections were acquired at 0.2 μm steps using a Plan Apo 100X/1.4 NA objective and appropriate fluorescence filters. Image analysis was performed in Fiji (ImageJ, NIH).

### Western blots

Cells were treated with 330 nM nocodazole or 10 μM STLC for 16-18 hrs before harvesting for Western blot analysis. Cells were washed with PBS and immediately lysed on ice for 30 min in fresh urea lysis buffer (50 mM Tris pH 7.5, 150 mM NaCl, 0.5% NP-40, 0.1% SDS, 6.5 M Urea, 1X Complete EDTA-free protease inhibitor cocktail (Roche), 1 mM PMSF). Cellular debris was removed by centrifugation. Protein concentrations in each sample were measured using either Bradford reagent (Bio-Rad) or Non-Interfering protein assay (G-Biosciences), and sample concentrations were normalized by addition of Laemmli sample buffer. Lysates were heated at 95°C for 5 min, separated by SDS-PAGE on a 10% acrylamide gel, and transferred to PVDF membrane (GE Healthcare or EMD Millipore).

For standard chemiluminescence, membrane was blocked for 1 hr in Blocking Buffer (2% milk in PBS + 0.05% Tween-20). Primary antibodies were diluted in 0.2% milk in PBS + 0.05% Tween-20 + 0.2% NaN3 and applied to the membrane overnight at 4°C or for 1-2 hrs at room temperature. HRP-conjugated secondary antibodies (GE Healthcare or Kindle Biosciences) were diluted in 0.2% milk in PBS + 0.05% Tween-20 and applied to the membrane for 1 hr at room temperature. After washing in PBS + 0.05% Tween-20, Clarity enhanced chemiluminescence substrate (Bio-Rad) was added to the membrane according to the manufacturer’s instructions. Membrane was imaged with a KwikQuant Imager (Kindle Biosciences) and analyzed using Adobe Photoshop.

For quantitative western, membrane was blocked for 1 hr in Intercept® Blocking Buffer (LI-COR). Primary antibodies were diluted in Intercept® Blocking Buffer + 0.05% Tween-20 and applied to the membrane for 1-2 hrs at room temperature. IRDye® 800CW Goat anti-mouse and/or IRDye® 680RD Goat anti-rabbit secondary antibodies (LI-COR) were diluted in Intercept® Blocking Buffer + 0.1% Tween-20 and applied to the membrane for 1 hr at room temperature. Membrane was imaged with the Odyssey® CLx Imager (LI-COR) and analyzed using LI-COR Image Studio software.

The following primary antibodies were used: C-terminus (aa 450-499) of human Cdc20 (C-term Ab, 1:500, Abcam, ab26483), N-terminus (aa 1-175) of human Cdc20 (N-term Ab, 1:300, Santa Cruz Biotechnology, sc-13162), antibody against acetylated M88-terminus of human Cdc20 (M88Ac Ab, 1:40,000, Cheeseman Lab), GAPDH (1:10,000, Cell Signaling Technology, #2118), GAPDH (1:20,000, Abcam, ab185059, HRP-conjugated), β-actin (1:20,000, Santa Cruz Biotechnology, sc-47778, HRP-conjugated).

### Lambda phosphatase treatment

HeLa cells were treated with 330 nM nocodazole overnight and mitotic cells were harvested by shake-off. Cells were washed with PBS and then lysed on ice for 45 min in HEPES/ Triton X-100 lysis buffer (20 mM HEPES pH 7.5, 150 mM NaCl, 1% Triton X-100, 5 mM MgCl2, 1X Complete ED-TA-free protease inhibitor cocktail (Roche), 1 mM PMSF). Cellular debris was removed by centrifugation and the resulting lysate was split into three parts. One sample was left untreated and the other two were supplemented with 1X Protein MetalloPhosphatase buffer (New England Biolabs) and 1 mM MnCl2 only or together with 1 μl Lambda Protein Phosphatase (New England Biolabs). After incubation at 30°C for 30 min, reactions were stopped by addition of 2X Laemmli sample buffer and heated at 95°C for 5 min. Samples were analyzed by SDS-PAGE and Western blot.

### Cell synchronization using single thymidine arrest

HeLa cells were first arrested in S phase using 2.5 mM thymidine (Sigma) for 22 hrs and then washed and released into medium without thymidine. Cells were collected at various time points after the single thymidine release. At 6 hrs post-release, 330 nM nocodazole was added to arrest cells in mitosis. Harvested cells were analyzed by SDS-PAGE and Western blot.

### Time-lapse experiments for mitotic timing

Cells were first seeded in 12-well polymer-bottomed plates (Cellvis, P12-1.5P) and treated as indicated in the figure legends. Cells were later moved to CO2-independent media (Gibco) supplemented with either 10% FBS or 15% FBS for HEC-6 derived cell lines, 100 U/mL penicillin and streptomycin, and 2 mM L-glutamine before imaging at 37°C. Phase contrast images were acquired on a Nikon eclipse microscope equipped with a charge-coupled device camera (Clara, Andor) or a sCMOS camera (ORCA-Fusion BT, Hamamatsu) using a Plan Fluor 20X/0.5 NA objective at either 2 min, 5 min, or 10 min intervals. Time-lapse movies were analyzed using Fiji (ImageJ, NIH), ilastik 32 and CellProfiler 33. Image brightness and contrast were first adjusted in Fiji. Pix-el-based classification was performed on individual images using ilastik and the resulting probability map images were processed by CellProfiler to identify and track mitotic cells. Each mitotic cell was then confirmed manually and the mitotic duration was determined as the time from cell rounding at mitotic entry to cell flattening after mitotic exit.

### Live-cell fluorescence imaging

For live-cell fluorescence imaging, cells were seeded into 8-well glass-bottomed chambers (Ibidi or Cellvis) and moved into CO2-independent media (Gibco) before imaging at 37°C. DNA was stained with 0.2 μg/ml Hoechst-33342 (Invitrogen). Cells were imaged directly or after 1 hr incubation with 330 nM nocodazole. Images were acquired on a DeltaVision Core deconvolution microscope (Applied Precision) equipped with a CoolSnap HQ2 charge-coupled device camera and deconvolved using the Softworx software. Z-sections were acquired at 0.5 μm steps using a Plan Apo 100X/1.4 NA objective and appropriate fluorescence filters. Image analysis was performed in Fiji (ImageJ, NIH).

### Mitotic index analysis

To determine the mitotic index from microscope images, cells were seeded in 12-well polymer-bottomed plates (Cellvis, P12-1.5P) and treated as indicated in the figure legends before fixing in PBS + 4% formaldehyde for 10 min at room temperature. After washing with PBS, cells were incubated in Abdil containing 0.5 μg/ml Hoechst-33342 (Invitrogen) for 30 min. Cells were washed with PBS and stored in Abdil until ready to image. Images were acquired on a Nikon eclipse microscope equipped with a charge-coupled device camera (Clara, Andor) using a Plan Fluor 20X/0.5 NA objective and appropriate fluorescence filters. Image analysis was performed in Fiji (ImageJ, NIH). The mitotic index was determined by scoring the number of mitotic cells with condensed DNA and dividing by the total number of cells.

For the analysis of the mitotic index using flow cytometry, cells were collected by incubation for 10 min in PBS + 5 mM EDTA, washed once in PBS, then fixed in PBS + 2% formaldehyde for 10 min at room temperature. After washing with PBS + 0.1% Triton X-100, cells were first immunostained for histone H3 phosphorylated at serine residue 10 (H3pS10 Ab, 1:3,000, Abcam, ab5176) diluted in Abdil, followed by AlexaFluor 647-conjugated secondary antibody (Jackson ImmunoResearch Laboratories). The proportion of GFP-positive single cells also staining positive for H3pS10 was determined on an LSR Fortessa (BD Biosciences) flow cytometer and analyzed with FlowJo software. High levels H3pS10 were used as a marker of mitosis. At least 550 GFP-positive cells were analyzed per condition.

### Evolutionary conservation analysis

Protein sequences of Cdc20 orthologs from the following tetrapod species were aligned using COBALT 34: Homo sapiens (Human, NP_001246.2), Macaca mulatta (Rhesus, NP_001248046.1), Mus musculus (Mouse, NP_075712.2), Canis lupus familiaris (Dog, XP_003639578.1), Loxodonta africana (Elephant, XP_023410383.1), Gallus gallus (Chicken, NP_001006536.1), and Xenopus tropicalis (Xenopus, NP_988945.1).

For the altORF conservation analysis, the DNA sequence of canonical open-reading frames (ORFs) from all mammalian species with orthologs to human CDC20 (ENSG00000117399) were obtained from Ensembl Release 103 35. For species with multiple orthologous transcripts, only one ORF sequence (the most conserved relative to human) was used. An ORF analysis was performed to identify start and stop codons in 2 forward frames and visualized for the first 300 nt.

### GFP immunoprecipitation and Mass Spectrometry

IP-MS experiments were performed as described previously 36. Eluted proteins from control HeLa cells with C-terminal mEGFP tag were digested with mass-spectrometry grade Lys-C and trypsin (Promega). Eluted proteins from a cell line lacking full-length Cdc20 (CDC20_M1-fs-M43, Extended Fig. 2A) to maximize the identification of alternative isoforms were digested with Lys-C alone. Both cell lines were enriched in mitosis using nocodazole only for control cells or double thymidine synchronization with nocodazole for the CDC20_M1-fs-M43 cell line.

Harvested cells were washed in PBS and resuspended 1:1 in 1X Lysis Buffer (50 mM HEPES pH 7.4, 1 mM EGTA, 1 mM MgCl2, 100 mM KCl, 10% glycerol) then drop frozen in liquid nitrogen. Cells were thawed after addition of an equal volume of 1.5X lysis buffer supplemented with 0.075% NP-40, 1X Complete EDTA-free protease inhibitor cocktail (Roche), 1 mM PMSF, 20 mM beta-glycerophosphate, and 0.4 mM sodium orthovanadate. Cells were lysed by sonication and cleared by centrifugation. The supernatant was mixed with Protein A beads coupled to rabbit anti-GFP antibody (Cheeseman lab) and rotated at 4°C for 1 hr. Beads were washed five times in Wash Buffer (50 mM HEPES pH 7.4, 1 mM EGTA, 1 mM MgCl2, 300 mM KCl, 10% glycerol, 0.05% NP-40, 1 mM DTT, 10 μg/mL leupeptin/pepstatin/chymostatin). After a final wash in Wash Buffer without detergent, bound protein was eluted with 100 mM glycine pH 2.6. Eluted proteins were precipitated by addition of 1/5th volume trichloroacetic acid at 4°C overnight. Precipitated proteins were reduced with TCEP, alkylated with iodoacetamide, and digested with mass-spectrometry grade Lys-C and trypsin or Lys-C alone (Promega). Digested peptides were separated by liquid chromatography and analyzed on an Orbitrap Elite mass spectrometer (Thermo Fisher). Data were analyzed using Proteome Discover Software (Thermo Fisher).

### M88Ac antibody generation

The M88Ac antibody was generated against a synthesized acetylated-peptide with the following amino acid sequence: Ac-MEVASFLLSC (New England Peptide, Covance). Serum from immunized rabbit was depleted against an acetylated spanning peptide (Ac-PHRSAAQMEVASFLC) and affinity-purified against the acetylated peptide.

### MTT viability assay

HeLa or M1-stop cells were seeded at a density of 2,000 cells per well in 96-well plates and subsequently cultured for 72 hrs in triplicate with increasing concentrations of the indicated anti-mitotic drug. After 72 hrs incubation, the medium was removed and the cells were stained with 2.5 mg/ ml MTT (3-(4,5-dimethylthiazol-2-yl)-2,5-diphenyltetrazolium bromide, Biosynth) in medium without serum for 3 hrs. Formazan crystals formed by the cells were then dissolved in 4 mM HCl and 0.1% NP-40 in isopropanol for 15 min. The absorbance was read at 570 nm on a Multiskan GO plate reader (Thermo Scientific), using 650 nm as reference wavelength.

Representative clonal cell lines expressing varying levels of the wild-type CDC20 cDNA from an inducible construct in the M1-stop mutant cell line were seeded at a density of 2,000 cells per well in 96-well plates and either cultured uninduced in triplicate or with CDC20(WT) induction for 43 hrs with 20 ng/ml. Cells were then cultured for 72 hrs with increasing concentrations of STLC and processed for MTT viability assay as described above.

HEC-6 cell lines expressing either mEGFP alone or CDC20(WT) or CDC20-ΔM1 were seeded in triplicate at a density of 1,500 cells per well in 96-well plates. Cells were transfected with either a non-targeting control pool for mEG-FP-expressing cells or Cdc20 siRNAs for cells expressing CDC20 constructs. siRNAs were applied as described above with volumes adjusted for 96-well plate format and transfection medium was changed after 6 hrs. At 32 hrs post-transfection, cells were then cultured for 92 hrs with increasing concentrations of STLC and processed for MTT viability assay as described above.

## Acknowledgements

The authors thank Kara McKinley for initial observations, Eric Spooner for assistance with the mass spectrometry analysis, George Bell for analyzing the altORF conservation, and Angelika Amon, Dave Bartel, and members of the Cheeseman lab for their critical input and suggestions. This work was supported by grants to IMC from the NIH/National Institute of General Medical Sciences (R35GM126930), National Science Foundation (2029868), the Moore Foundation, the American Cancer Society (Theory Lab Collaborative Grant (TLC-20-117-01-TLC)), a Pilot award from the Global Consortium for Reproductive Longevity and Equity (GCRLE-1220), and a Hope Funds for Cancer Research fellowship (HFCR-18-03-02) to MJT.

## Author Contributions

Conceptualization – MJT, IMC; Methodology – MJT; Validation – MJT; Investigation – MJT; Writing - Original Draft Preparation – MJT, IMC; Writing – Review & Editing – MJT, IMC; Visualization – MJT; Supervision: IMC; Funding Acquisition: IMC, MJT

## Additional Information

The data that support the findings of this study are available from the corresponding author upon reasonable request. Correspondence and requests for materials should be addressed to Iain Cheeseman.

## Supplemental Information

**Extended Data Figure 1.**
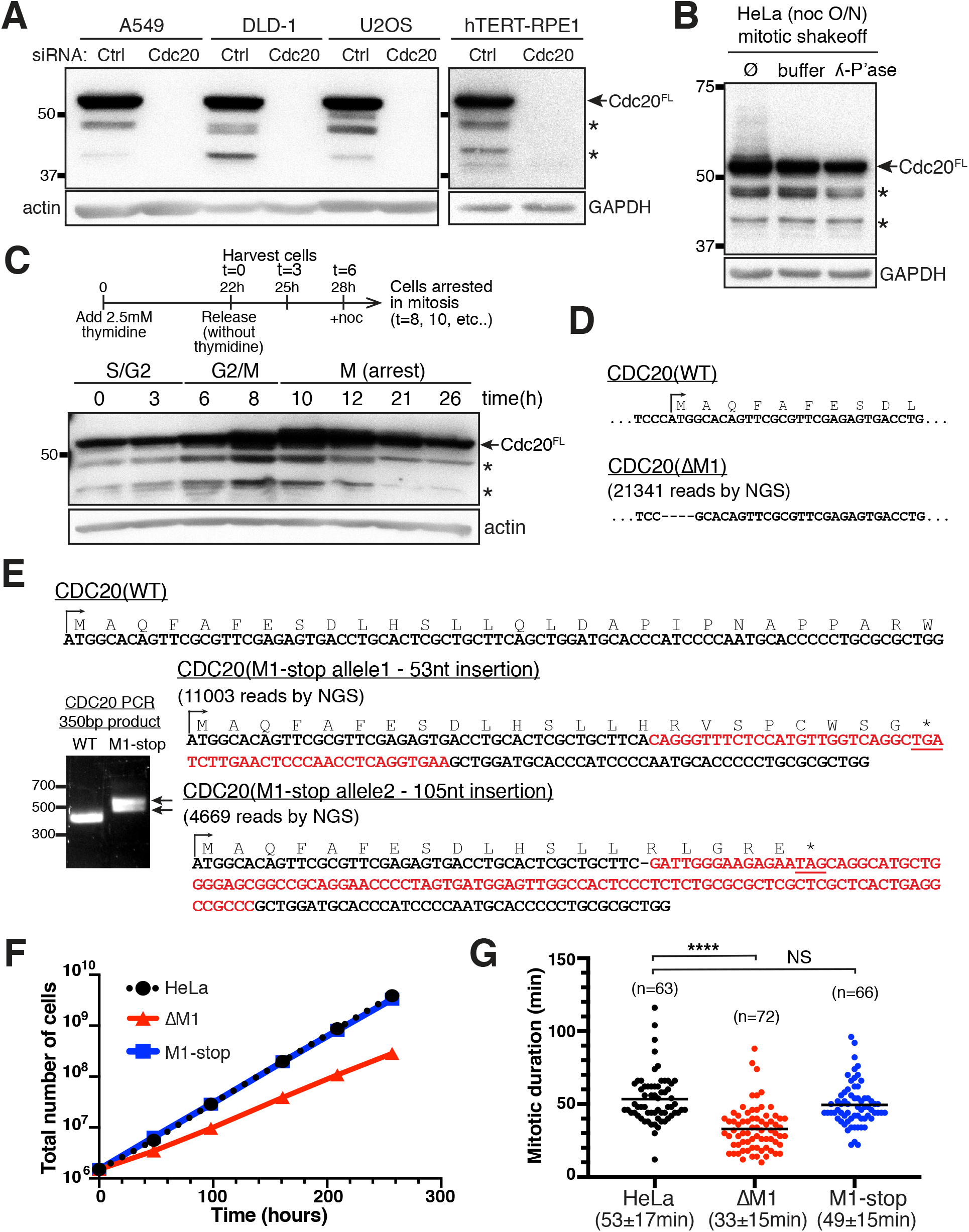
Human cells express alternative isoforms of Cdc20. Related to Figure 1. (**A**) Western blot showing multiple human cancer cell lines (A549, DLD-1, and U2OS) and the non-transformed hTERT RPE-1 cells treated with control or Cdc20 siRNAs and probed with antibodies recognizing the C-terminus of human Cdc20 (aa 450-499). β-actin or GAPDH was used as loading control. (**B**) Western blot showing mitotic HeLa cells collected by shake-off after overnight treatment with 330 nM nocodazole. Lysates alone, with buffer only, or with buffer and lambda phosphatase treatment were probed using antibodies recognizing the C-terminus of human Cdc20 (aa 450-499). GAPDH was used as loading control. (**C**) Schematic illustrating cell synchronization approach using a single thymidine arrest in S phase and subsequent release into the cell cycle. Addition of 330 nM nocodazole leads to a prolonged mitotic arrest. Lysates of HeLa cells harvested at the indicated time points after single thymidine release were separated by SDS-PAGE and endogenous Cdc20 protein was detected using antibodies recognizing the C-terminus of human Cdc20 (aa 450-499). β-actin was used as loading control. (**D**) Sequence information for the homozygous ΔM1 mutant cell line lacking the canonical M1 ATG start codon. The DNA sequence of the genomic locus was determined by next-generation sequencing. (**E**) Sequence information for the M1-stop mutant cell line showing insertions of 53 nt and 105 nt respectively after the L14 residue. Underlined are premature stop codons that are in-frame with the M1 ATG start codon for both *CDC20* alleles. The DNA sequence of the genomic locus was determined by next-generation sequencing. (**F**) Growth curves of control HeLa compared to the ΔM1 and M1-stop mutant cell lines. (**G**) Graph showing mitotic duration for control HeLa compared to the ΔM1 and M1-stop mutant cell lines. Each point represents a single cell. The bars correspond to the mean. Indicated are the mean mitotic duration ± standard deviation across two experimental replicates. Statistics from Student’s two-sample t-Test with two-tailed distribution (**** = p < 0.0001, NS = not significant).

**Extended Data Figure 2.**
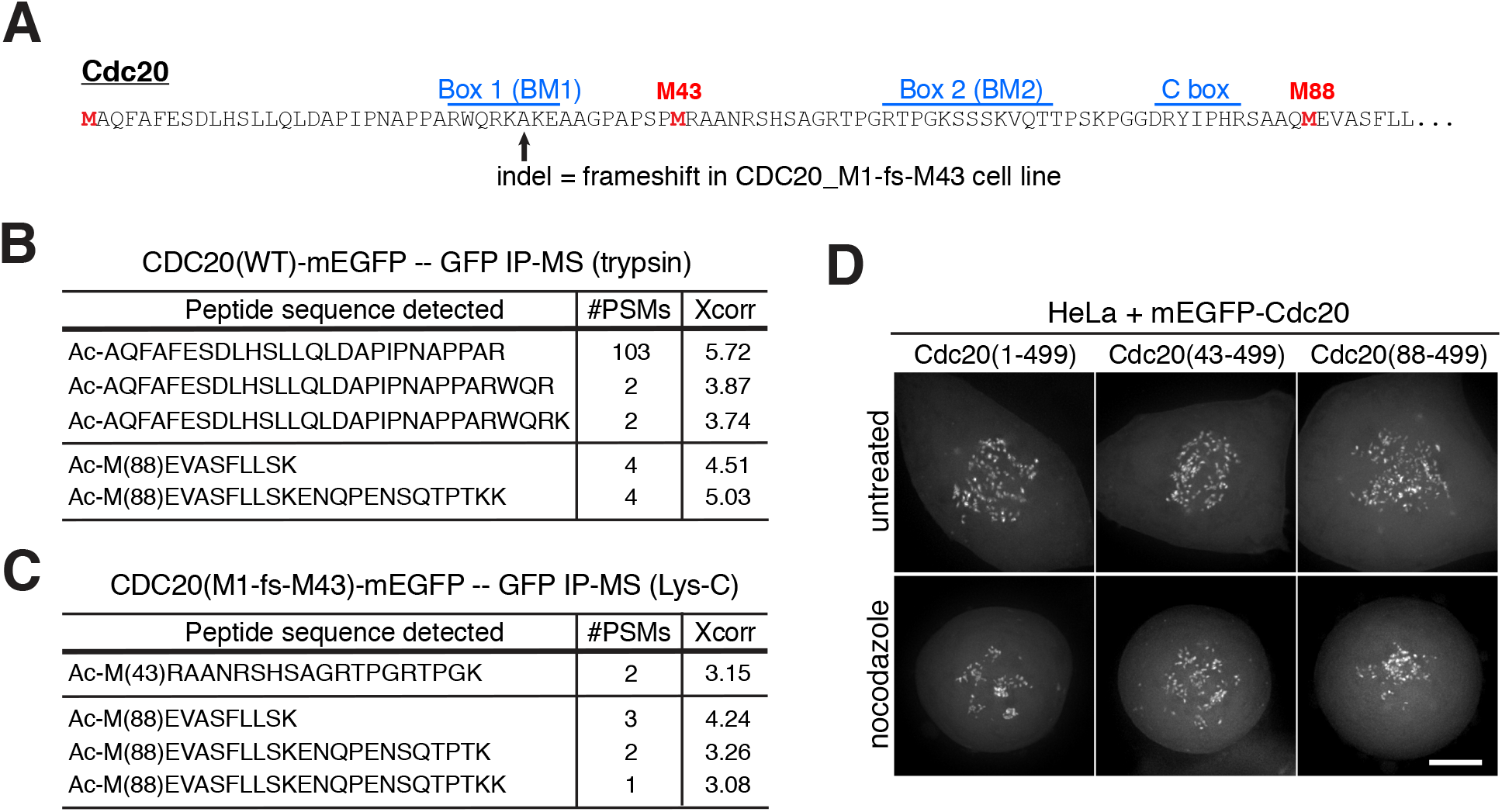
Cdc20 isoforms are produced by alternative translation initiation at downstream in-frame start codons. Related to Figure 1. (**A**) Protein sequence of the N-terminal region of human Cdc20 with conserved motifs indicated. Arrow indicates the location of indel mutations in the *CDC20*_M1-fs-M43 cell line. (**B**) Cdc20 tryptic peptides with N-terminal acetylation indicative of translation initiation were identified following immunoprecipitation-mass spectrometry of mitotically-enriched samples using an endogenous C-terminal mEGFP tag. The identified peptide sequence, number of peptide-spectrum matches (#PSMs), and the cross-correlation (Xcorr) value from the SEQUEST search are indicated. (**C**) Cdc20 peptides as in (B), except using the endopeptidase LysC and isolated using a C-terminal mEGFP tag in a cell line lacking the full-length Cdc20 protein (*CDC20*_M1-fs-M43). (**D**) Representative live-cell fluorescence microscopy images of untreated or nocodazole-treated HeLa cells expressing the indicated N-terminal mEGFP-Cdc20 fusions with 5 ng/ml doxycycline. Images show maximum intensity projections of deconvolved Z-stacks of selected mitotic cells. Images were scaled individually to highlight kinetochore localization. Scale bar, 5 μm.

**Extended Data Figure 3.**
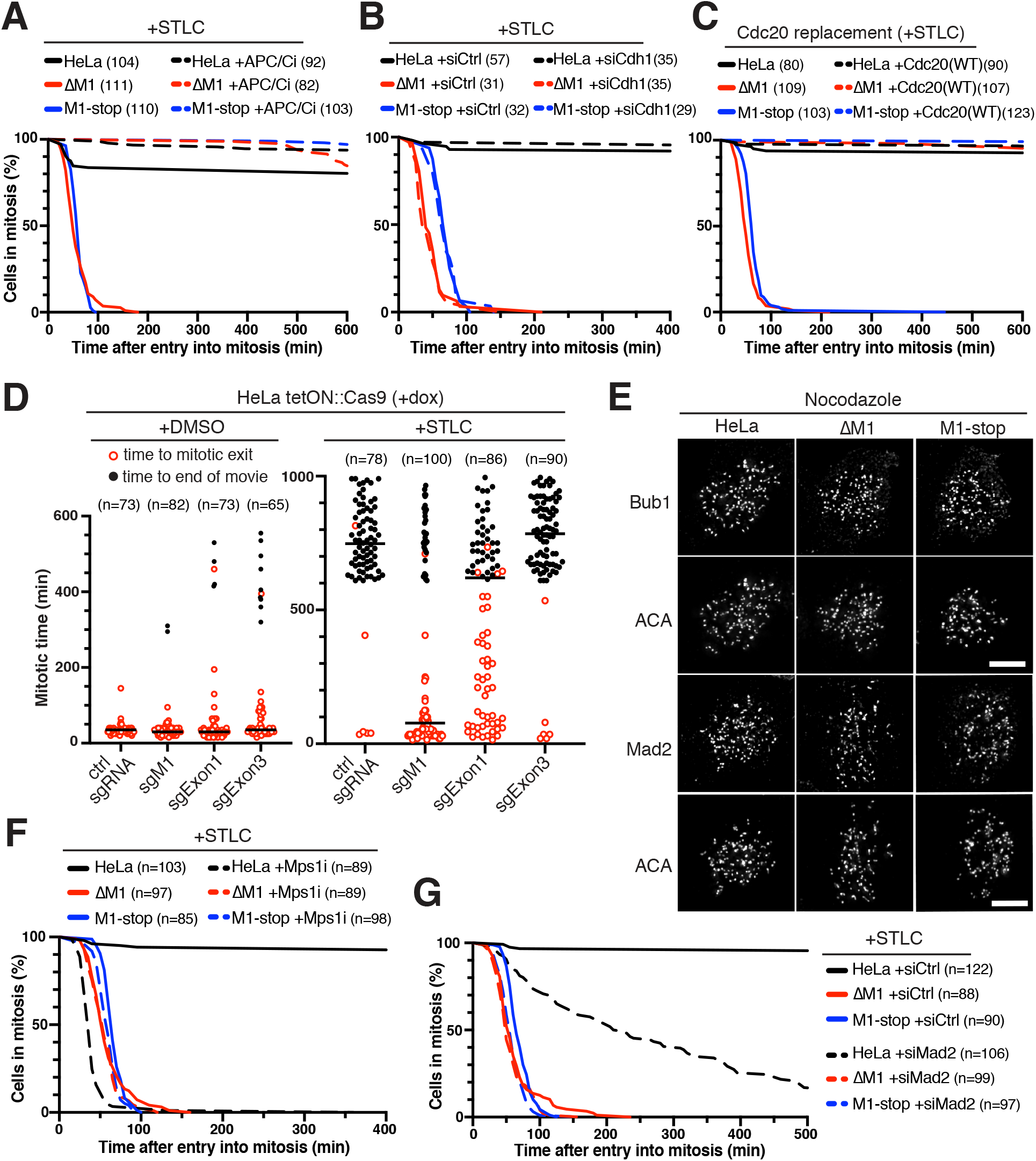
Truncated Cdc20 isoforms are inefficient targets of the SAC and promote mitotic slippage. Related to Figure 2. (**A**) Cumulative frequency distribution showing the fraction of mitotic cells over time post-mitotic entry for HeLa, ΔM1, and M1-stop cells treated with 10 μM STLC alone or in combination with the APC/C-inhibitor proTAME (12 μM). (**B**) Cumulative frequency distribution for the fraction of cells in mitosis at the indicated time after entry into mitosis (mitotic arrest duration) for HeLa, ΔM1, and M1-stop cells treated with 10 μM STLC and 100nM of either control siRNAs or Cdh1 siRNAs. (**C**) Cumulative frequency distribution for the fraction of cells in mitosis in the presence of 10 μM STLC for HeLa, ΔM1, and M1-stop cells expressing endogenous Cdc20 protein or upon Cdc20 replacement with ectopic wild-type *CDC20* cDNA. (**D**) Mitotic duration of individual HeLa cells expressing doxycycline-inducible Cas9 and sgRNAs recognizing different regions within the *CDC20* gene. Unperturbed mitotic progression or mitotic arrest behavior were monitored upon treatment with DMSO or 10 μM STLC respectively. Cells entering mitosis in the first 350-400 min of time lapse experiments were included in the analyses. Open red circles indicate cells that exit mitosis. Closed black circles indicate cells that remained arrested in mitosis till the end of the time lapse. Bars correspond to the median. (**E**) Representative immunofluorescence images of Bub1 or Mad2 localization to kinetochores immuno-stained with anti-centromere antibodies (ACA). Images are maximum intensity projections of deconvolved Z-stacks of selected mitotic cells from control HeLa or mutant ΔM1 or M1-stop cell lines treated with nocodazole. Images were scaled individually to highlight kinetochore localization. Scale bar, 5 μm. (**F**) Cumulative frequency distribution of the fraction of mitotic cells over time post-mitotic entry for HeLa, ΔM1, and M1-stop cells treated with 10 μM STLC alone or in combination with the Mps1-inhibitor AZ3146 (4 μM). (**G**) Cumulative frequency distribution showing the fraction of mitotic cells over time post-mitotic entry for HeLa, ΔM1, and M1-stop cells treated with 10 μM STLC and either control siRNAs or Mad2 siRNAs.

**Extended Data Figure 4.**
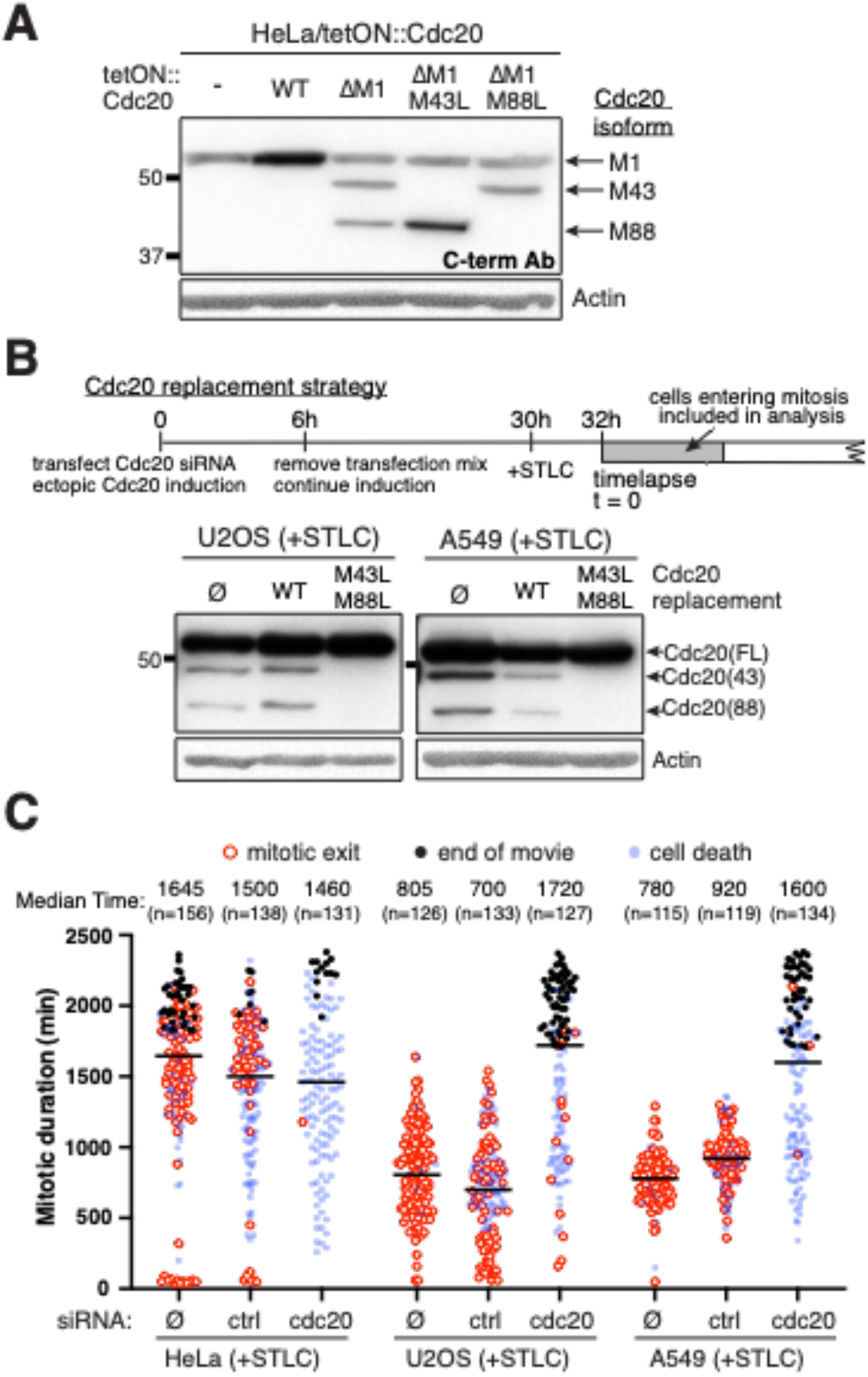
Cdc20 translational isoforms modulate mitotic arrest duration. Related to Figure 2. (**A**) Western blot showing HeLa cells alone or treated with 50 ng/ml doxycycline to express the indicated doxycycline-inducible *CDC20* constructs. Cdc20 protein was detected using antibodies recognizing the C-terminus of human Cdc20 (aa 450-499). β-actin was used as loading control. (**B**) Top, schematic illustrating the Cdc20 replacement strategy with siRNA-resistant 5’ UTR-*CDC20* cDNA constructs from a doxycycline-inducible promoter combined with depletion of endogenous Cdc20 protein by siRNA treatment. Bottom, Western blot of mitotically-enriched U2OS or A549 cells expressing endogenous Cdc20 protein or upon Cdc20 replacement with either wild-type *CDC20* cDNA or a Cdc20 M43L M88L mutant construct. Cells were enriched in mitosis with 10 μM STLC for 18 hrs. Cdc20 protein was detected using antibodies recognizing the human Cdc20 C-terminus (aa 450-499). β-actin was used as loading control. (**C**) Mitotic arrest duration of individual HeLa, U2OS, or A549 cells treated with 10 μM STLC alone or with a combination of STLC and siRNA treatment (either control siRNAs or Cdc20 siRNAs). Cells entering mitosis in the first 600 min (HeLa) or 700 min (U2OS/A549) of time lapse experiments were included in analyses. Open red circles indicate cells that exit mitosis. Closed black circles indicate cells that remained arrested in mitosis till the end of the time lapse. Blue circles indicate cells that die in mitosis. Bars correspond to median. Indicated are the median mitotic duration times across two experimental replicates.

**Extended Figure 5.**
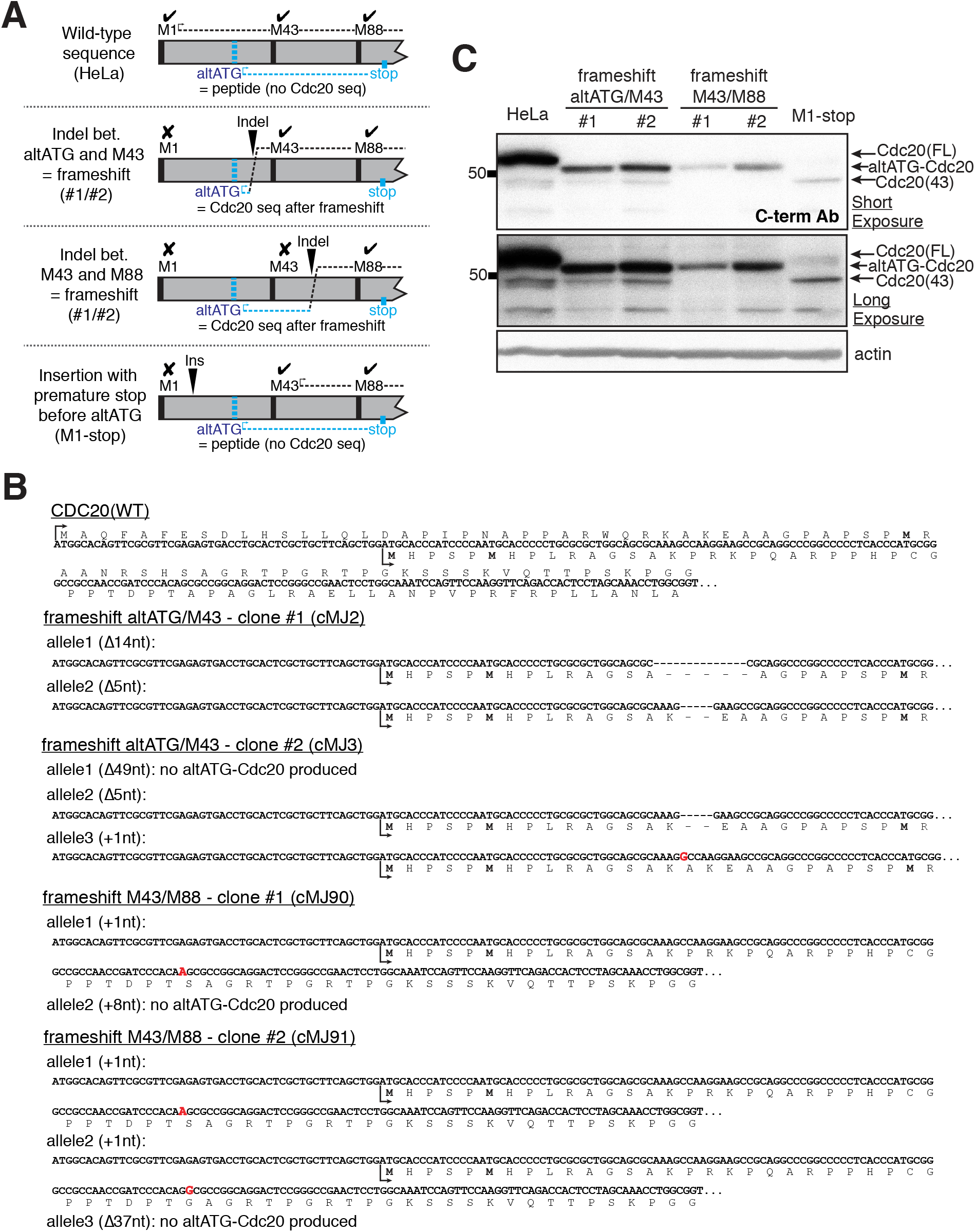
Translation initiation at alternative out-of-frame start codons in HeLa cells. Related to Figure 3. (**A**) Schematic illustrating the strategy to assess whether translation initiation occurs at the alternative out-of-frame start codons. Cdc20 protein indicated in black; altORF peptide indicated in cyan. Cell lines with indel mutations after the alternative start codons at the endogenous locus in all *CDC20* alleles disrupt translation of the full-length Cdc20 protein. Some indel mutations resulted in a frame shift that would be predicted to connect the altORF peptide with amino acid sequences encoding downstream regions of Cdc20. If the altORF is translated, this would produce a chimeric protein (altATG-Cdc20) that is shorter than full-length Cdc20, but is detectable with antibodies against the Cdc20 C-terminus. In contrast, insertions upstream of the alternative out-of-frame start codons, such as those in the M1-stop mutant, should only abrogated expression of the full-length Cdc20 protein without generating a chimeric altATG-Cdc20 protein. (**B**) Analysis of wild-type human *CDC20* nucleic acid sequence reveals two alternative out-of-frame start codons between Met1 and Met43. The amino acid sequence of the predicted alternative open reading frame (altORF) is indicated below the nucleic acid sequence, with the methionines bolded. Sequence information is shown for representative clones with indel mutations where at least one *CDC20* allele results in a frame shift that connects the altORF peptide with amino acid sequences encoding downstream regions of Cdc20. The DNA sequence of the genomic locus was determined by next-generation sequencing. Insertions are highlighted in red. When present, the amino acid sequence of the resulting altATG-Cdc20 peptide produced is shown. (**C**) Western blot showing mitotically-enriched control HeLa, M1-stop mutant, and representative clones with indel mutations where at least one *CDC20* allele resulted in a frame shift that connects the altORF peptide with amino acid sequences encoding downstream regions of Cdc20. Endogenous Cdc20 protein was detected using antibodies recognizing the C-terminus of human Cdc20 (aa 450-499). β-actin was used as loading control.

**Extended Data Figure 6.**
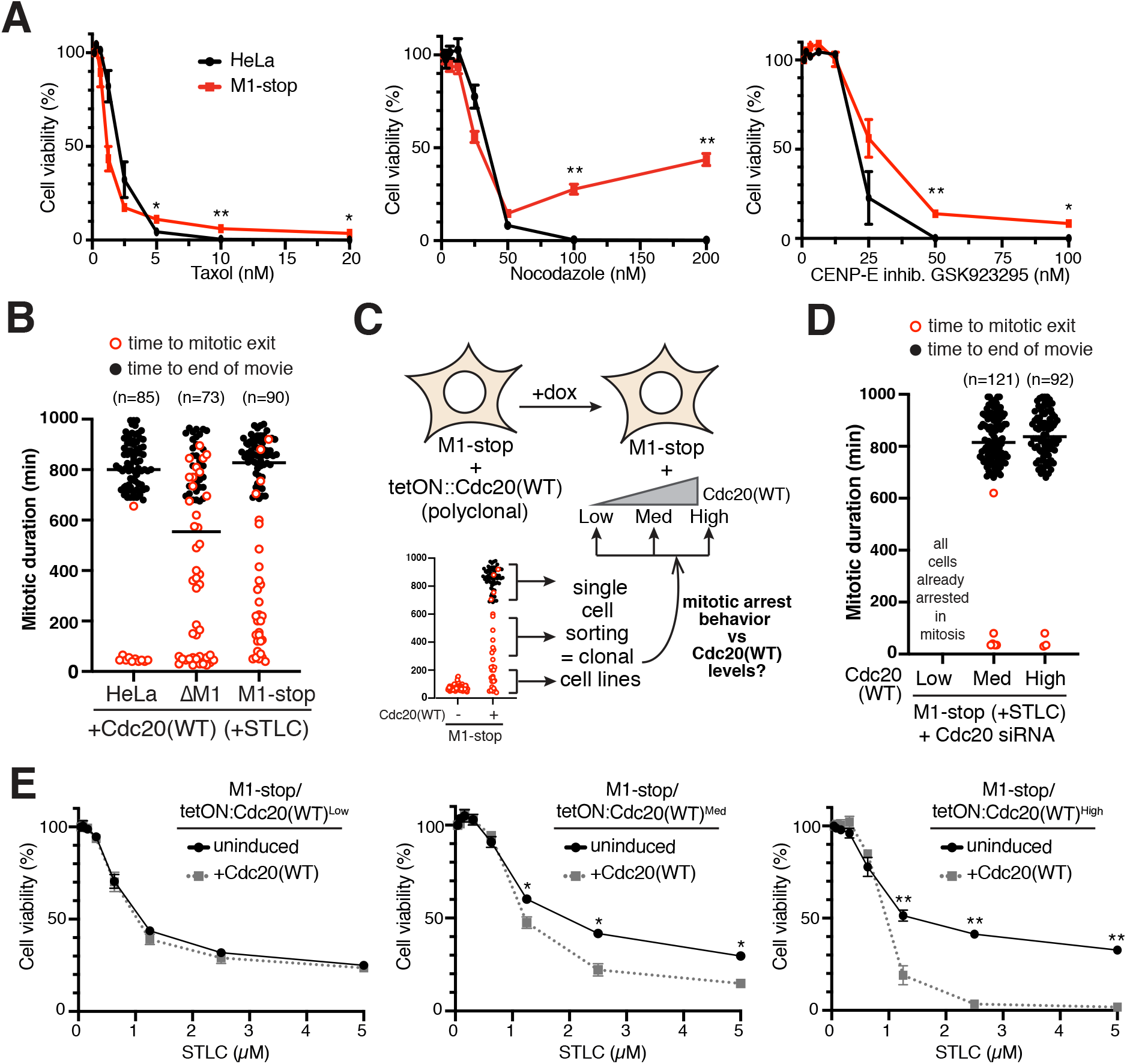
Cdc20 translational isoform levels alter cancer cell anti-mitotic drug sensitivity. Related to Figure 4. (**A**) Sensitivity of HeLa or M1-stop cells to increasing concentrations of Taxol, Nocodazole, or the CENP-E inhibitor GSK923295. Cell viability was determined by MTT assay in triplicate following 72 h drug treatment. Error bars indicate SEM of three (Nocodazole) or four (Taxol, GSK923295) experimental replicates. Statistics from Student’s two-sample t-Test with two-tailed distribution comparing HeLa and M1-stop cell viabilities per drug concentration (* = p < 0.05, ** = p < 0.01). (**B**) Mitotic arrest duration in the presence of 10 μM STLC for individual HeLa, ΔM1, or M1-stop cells expressing the wild-type *CDC20* cDNA. Open red circles indicate cells that exit mitosis. Closed black circles indicate cells that remained arrested in mitosis till the end of the time lapse. Bars correspond to the median. (**C**) Schematic illustrating the approach to isolate clonal cell lines from the polyclonal M1-stop mutant expressing the doxycycline-inducible wild-type *CDC20* construct. Multiple clones were analyzed to assess the correlation between the mitotic arrest behavior of a given clone and the expression level of the integrated doxycycline-inducible *CDC20* construct (see text for details). (**D**) Similar mitotic arrest duration as in (B) except for representative clones of M1-stop mutant with low, medium, or high expression of the doxycycline-inducible wild-type *CDC20* construct. Cells were treated with Cdc20 siRNAs to deplete endogenous truncated Cdc20 isoforms. Low Cdc20 expression from the inducible *CDC20* construct fails to support mitotic progression even before addition of STLC. (**E**) Similar sensitivity assay as in (A) except for STLC for representative clones of M1-stop mutant with low, medium, or high expression of the doxycycline-inducible wild-type *CDC20* construct without induction or induced with 20 ng/μl doxycycline. Error bars indicate SEM of three experimental replicates.

**Extended Data Table 1.**
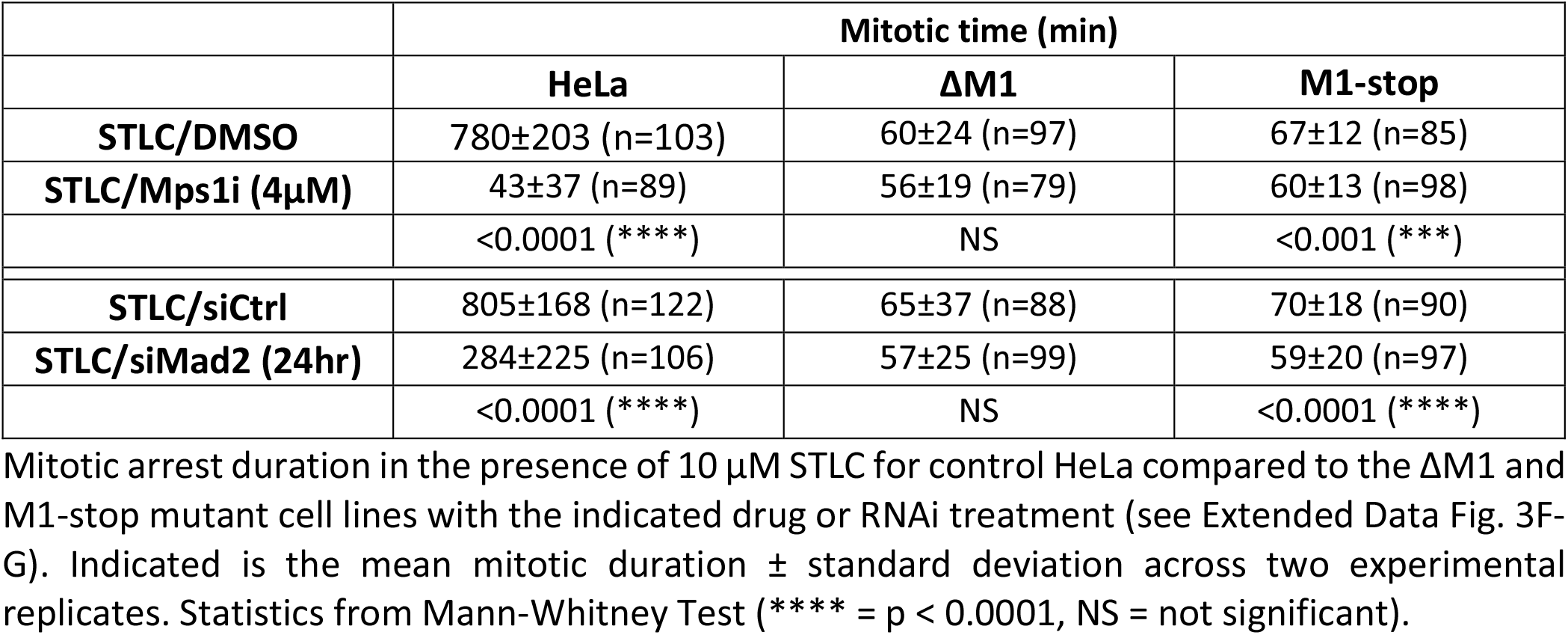
The SAC is defective in ΔM1 and M1-stop mutant cell lines. Related to Extended Data Figure 3.

|  | Mitotic time (min) |  |  |
| --- | --- | --- | --- |
| | HeLa | ΔM1 | M1-stop |
| **STLC/DMSO** | 780±203 (n=103) | 60±24 (n=97) | 67±12 (n=85) |
| **STLC/Mps1i (4μM)** | 43±37 (n=89) | 56±19 (n=79) | 60±13 (n=98) |
|  | <0.0001 (****) | NS | <0.001 (***) |
| **STLC/siCtrl** | 805±168 (n=122) | 65±37 (n=88) | 70±18 (n=90) |
| **STLC/siMad2 (24hr)** | 284±225 (n=106) | 57±25 (n=99) | 59±20 (n=97) |
|  | <0.0001 (****) | NS | <0.0001 (****) |

**Extended Data Table 2.**
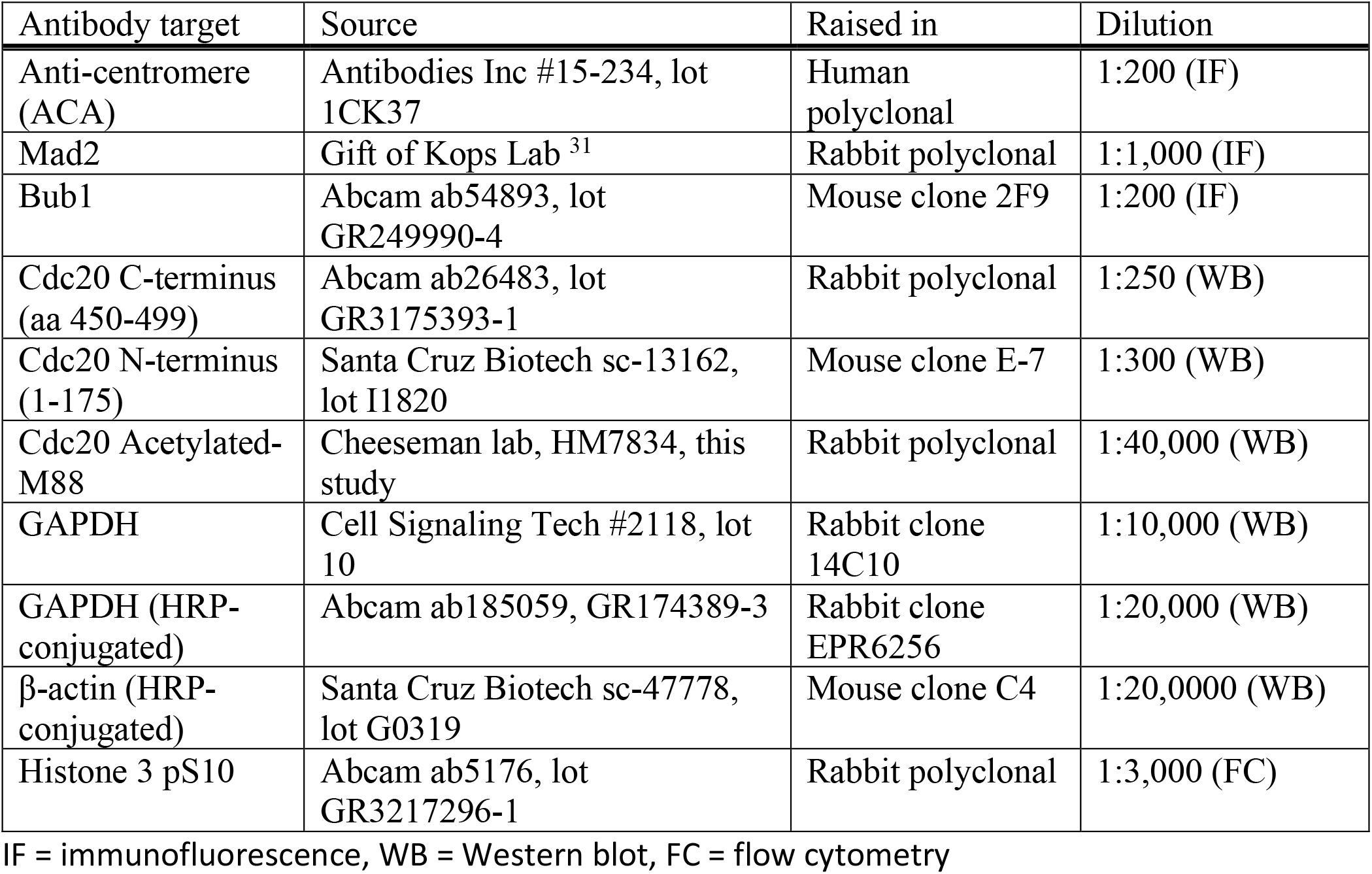
Antibodies used in this study.

| Antibody target | Source | Raised in | Dilution |
| --- | --- | --- | --- |
| Anti-centromere (ACA) | Antibodies Inc #15-234, lot 1CK37 | Human polyclonal | 1:200 (IF) |
| Mad2 | Gift of Kops Lab <sup>31</sup> | Rabbit polyclonal | 1:1,000 (IF) |
| Bub1 | Abcam ab54893, lot GR249990-4 | Mouse clone 2F9 | 1:200 (IF) |
| Cdc20 C-terminus (aa 450-499) | Abcam ab26483, lot GR3175393-1 | Rabbit polyclonal | 1:250 (WB) |
| Cdc20 N-terminus (1-175) | Santa Cruz Biotech sc-13162, lot I1820 | Mouse clone E-7 | 1:300 (WB) |
| Cdc20 Acetylated-M88 | Cheeseman lab, HM7834, this study | Rabbit polyclonal | 1:40,000 (WB) |
| GAPDH | Cell Signaling Tech #2118, lot 10 | Rabbit clone 14C10 | 1:10,000 (WB) |
| GAPDH (HRP-conjugated) | Abcam ab185059, GR174389-3 | Rabbit clone EPR6256 | 1:20,000 (WB) |
| β-actin (HRP-conjugated) | Santa Cruz Biotech sc-47778, lot G0319 | Mouse clone C4 | 1:20,000 (WB) |
| Histone 3 pS10 | Abcam ab5176, lot GR3217296-1 | Rabbit polyclonal | 1:3,000 (FC) |

IF = immunofluorescence, WB = Western blot, FC = flow cytometry

**Extended Data Table 3.** Cell lines used in this study.

| Name | Description | Expression | Source |
| --- | --- | --- | --- |
| A549 | Parental | NA | Lab stock |
| HeLa | Parental | NA | Lab stock |
| hTERT-RPE1 | Parental | NA | Lab stock |
| U2OS | Parental | NA | Lab stock |
| HEC-6 | Parental | NA | JCRB Cell Bank |
| cTT20 | HeLa tetON::spCas9 | Dox-inducible Cas9 | <sup>5</sup> |
| cKC443 | cTT20 / Ctrl sgRNA | Dox-inducible Cas9 | Lentiviral transduction of pKC328 |
| cMJ2 | cTT20_frameshift altATG/M43 - clone #1 | NA | Lentiviral transduction of pKMKO D4.1 + dox induction; clonal |
| cMJ3 | cTT20_frameshift altATG/M43 - clone #2 | NA | Lentiviral transduction of pKMKO D4.1 + dox induction; clonal |
| cMJ5 | cTT20 / sgExon3 | Dox-inducible Cas9 | Lentiviral transduction of pMJ7 |
| cMJ15 | cTT20_cdc20-mEGFP (heterozygous) | Dox-inducible Cas9 | Transient transfection of pMJ19 + pMJ27; clonal |
| cMJ35 | cMJ15_frameshift altATG/M43 | NA | Lentiviral transduction of pKMKO D4.1 + dox induction; clonal |
| cMJ46 | cMJ5 / Cdc20(1-499)_IRES2_EGFP | Dox-inducible Cas9/Constitutive retroviral | Retroviral transduction of pMJ39 |
| cMJ47 | cMJ5 / Cdc20(88-499)_IRES2_EGFP | Dox-inducible Cas9/Constitutive retroviral | Retroviral transduction of pMJ41 |
| cMJ90 | cTT20_frameshift M43/M88 - clone #1 | NA | Lentiviral transduction of pMJ32 + dox induction; clonal |
| cMJ91 | cTT20_frameshift M43/M88 - clone #2 | NA | Lentiviral transduction of pMJ32 + dox induction; clonal |
| cMJ94 | cMJ5 / Cdc20(43-499)_IRES2_EGFP | Dox-inducible Cas9/Constitutive retroviral | Retroviral transduction of pMJ40 |
| cMJ96 | cMJ5 / empty_IRES2_EGFP | Dox-inducible Cas9/Constitutive retroviral | Retroviral transduction of pMJ31 |
| cMJ159 | cTT20 / sgExon1 | Dox-inducible Cas9 | Lentiviral transduction of pMJ92 |
| cMJ200 | HeLa_M1-stop | NA | Transient transfection of pMJ94; clonal |
| cMJ222 | HeLa_ΔM1 | NA | Transient transfection of pMJ118; clonal |
| cMJ226 | cTT20 / sgM1 | Dox-inducible Cas9 | Retroviral transduction of pMJ141 |
| cMJ230 | cMJ200 /<br>kozak_cdc20(WT)_3UTR | Dox-inducible<br>AAVS1 | AAVS1 Safe-harbor insertion of<br>pMJ143 |
| cMJ231 | cMJ200 /<br>5UTR_cdc20(WT)_3UTR | Dox-inducible<br>AAVS1 | AAVS1 Safe-harbor insertion of<br>pMJ144 |
| cMJ233 | cMJ200 /<br>kozak_cdc20(M43L)_3UT<br>R | Dox-inducible<br>AAVS1 | AAVS1 Safe-harbor insertion of<br>pMJ146 |
| cMJ234 | cMJ200 /<br>kozak_cdc20(M88L)_3UT<br>R | Dox-inducible<br>AAVS1 | AAVS1 Safe-harbor insertion of<br>pMJ147 |
| cMJ236 | cMJ200 /<br>5UTR_cdc20(altATGmutx<br>2)_3UTR | Dox-inducible<br>AAVS1 | AAVS1 Safe-harbor insertion of<br>pMJ149 |
| cMJ241 | HeLa /<br>kozak_cdc20(WT)_3UTR | Dox-inducible<br>AAVS1 | AAVS1 Safe-harbor insertion of<br>pMJ143 |
| cMJ242 | cMJ222 /<br>kozak_cdc20(WT)_3UTR | Dox-inducible<br>AAVS1 | AAVS1 Safe-harbor insertion of<br>pMJ143 |
| cMJ253 | HeLa /<br>5UTR_cdc20(WT)_3UTR | Dox-inducible<br>AAVS1 | AAVS1 Safe-harbor insertion of<br>pMJ144 |
| cMJ256 | HeLa /<br>5UTR_cdc20(altATGmutx<br>2)_3UTR | Dox-inducible<br>AAVS1 | AAVS1 Safe-harbor insertion of<br>pMJ149 |
| cMJ261 | cMJ200 /<br>kozak_cdc20(WT-low)_3UTR | Dox-inducible<br>AAVS1 | Single-cell sort cMJ230; clonal |
| cMJ263 | cMJ200 /<br>kozak_cdc20(WT-med)_3UTR | Dox-inducible<br>AAVS1 | Single-cell sort cMJ230; clonal |
| cMJ264 | cMJ200 /<br>kozak_cdc20(WT-high)_3UTR | Dox-inducible<br>AAVS1 | Single-cell sort cMJ230; clonal |
| cMJ268 | HeLa / mEGFP-cdc20(1-499) | Dox-inducible<br>AAVS1 | AAVS1 Safe-harbor insertion of<br>pMJ160 |
| cMJ269 | HeLa / mEGFP-cdc20(43-499) | Dox-inducible<br>AAVS1 | AAVS1 Safe-harbor insertion of<br>pMJ161 |
| cMJ270 | HeLa / mEGFP-cdc20(88-499) | Dox-inducible<br>AAVS1 | AAVS1 Safe-harbor insertion of<br>pMJ162 |
| cMJ274 | HeLa /<br>kozak_cdc20(ΔM1)_3UT<br>R | Dox-inducible<br>AAVS1 | AAVS1 Safe-harbor insertion of<br>pMJ163 |
| cMJ275 | HeLa / kozak_cdc20(ΔM1<br>M43L)_3UTR | Dox-inducible<br>AAVS1 | AAVS1 Safe-harbor insertion of<br>pMJ164 |
| cMJ276 | HeLa / kozak_cdc20(ΔM1 M88L)_3UTR | Dox-inducible AAVS1 | AAVS1 Safe-harbor insertion of pMJ165 |
| cMJ278 | cMJ200 / kozak_cdc20(ΔM1)_3UTR | Dox-inducible AAVS1 | AAVS1 Safe-harbor insertion of pMJ163 |
| cMJ279 | cMJ200 / kozak_cdc20(ΔM1 M43L)_3UTR | Dox-inducible AAVS1 | AAVS1 Safe-harbor insertion of pMJ164 |
| cMJ280 | cMJ200 / kozak_cdc20(ΔM1 M88L)_3UTR | Dox-inducible AAVS1 | AAVS1 Safe-harbor insertion of pMJ165 |
| cMJ281 | cMJ200 / kozak_cdc20(ΔM1 M43L M88L)_3UTR | Dox-inducible AAVS1 | AAVS1 Safe-harbor insertion of pMJ166 |
| cMJ285 | cMJ200 / 5UTR_cdc20(addATG)_3UTR | Dox-inducible AAVS1 | AAVS1 Safe-harbor insertion of pMJ175 |
| cMJ288 | HeLa / 5UTR_cdc20(M43L M88L)_3UTR | Dox-inducible AAVS1 | AAVS1 Safe-harbor insertion of pMJ176 |
| cMJ290 | U2OS / 5UTR_cdc20(WT)_3UTR | Dox-inducible AAVS1 | AAVS1 Safe-harbor insertion of pMJ144 |
| cMJ292 | U2OS / 5UTR_cdc20(M43L M88L)_3UTR | Dox-inducible AAVS1 | AAVS1 Safe-harbor insertion of pMJ176 |
| cMJ294 | A549 / 5UTR_cdc20(WT)_3UTR | Dox-inducible AAVS1 | AAVS1 Safe-harbor insertion of pMJ144 |
| cMJ296 | A549 / 5UTR_cdc20(M43L M88L)_3UTR | Dox-inducible AAVS1 | AAVS1 Safe-harbor insertion of pMJ176 |
| cMJ321 | cMJ200 / 5UTR_cdc20(optimal-Kozak)_3UTR | Dox-inducible AAVS1 | AAVS1 Safe-harbor insertion of pMJ186 |
| cMJ322 | cMJ200 / 5UTR_cdc20(anti-Kozak)_3UTR | Dox-inducible AAVS1 | AAVS1 Safe-harbor insertion of pMJ187 |
| cMJ348 | HEC-6 / mEGFP | Constitutive | Lentiviral transduction of pMJ197 |
| cMJ349 | HEC-6 / kozak_cdc20(WT) | Constitutive | Lentiviral transduction of pMJ206 |
| cMJ351 | HEC-6 / kozak_cdc20(ΔM1) | Constitutive | Lentiviral transduction of pMJ208 |

